# Breast cancer cells promote osteoclast differentiation in an MRTF-dependent paracrine manner

**DOI:** 10.1101/2023.12.06.570453

**Authors:** Pooja Chawla, Ishani Sharma, David Gau, Ian Eder, Fangyuan Chen, Virginia Yu, Niharika Welling, David Boone, Juan Taboas, Adrian V. Lee, Adriana Larregina, Deborah L. Galson, Partha Roy

## Abstract

Bone is a frequent site for breast cancer metastasis. The vast majority of breast cancer-associated metastasis is osteolytic in nature, and RANKL (receptor activator for nuclear factor κB)-induced differentiation of bone marrow-derived macrophages (BMDMs) to osteoclasts (OCLs) is a key requirement for osteolytic metastatic growth of cancer cells. In this study, we demonstrate that Myocardin-related transcription factor (MRTF) in breast cancer cells plays an important role in paracrine modulation of RANKL-induced osteoclast differentiation. This is partly attributed to MRTF’s critical role in maintaining the basal cellular expression of connective tissue growth factor (CTGF), findings that align with a strong positive correlation between CTGF expression and MRTF-A gene signature in the human disease context. Luminex analyses reveal that MRTF depletion in breast cancer cells has a broad impact on OCL-regulatory cell-secreted factors that extend beyond CTGF. Experimental metastasis studies demonstrate that MRTF depletion diminishes OCL abundance and bone colonization breast cancer cells *in vivo*, suggesting that MRTF inhibition could be an effective strategy to diminish OCL formation and skeletal involvement in breast cancer. In summary, this study highlights a novel tumor-extrinsic function of MRTF relevant to breast cancer metastasis.

**SIGNIFICANCE STATEMENT:**

- MRTF, a transcriptional coactivator of SRF, is known to promote breast cancer progression through its tumor-cell-intrinsic function. Whether and how MRTF activity in tumor cells modulates other types of cells in the tumor microenvironment are not clearly understood.
- This study uncovers a novel tumor-cell-extrinsic function of MRTF in breast cancer cells in promoting osteoclast differentiation partly through CTGF regulation, and further demonstrates MRTF’s requirement for bone colonization of breast cancer cells in vivo.
- Our studies suggest that MRTF inhibition could be an effective strategy to diminish osteoclast formation and skeletal involvement in metastatic breast cancer.

## INTRODUCTION

Breast cancer is one of the leading causes of cancer-related death in women. The vast majority of breast cancer-related deaths are due to the metastatic spread of tumor cells to distant organs. Bone is a frequent site of breast cancer metastasis with the median survival of patients after tumor cells have metastasized to the bone being 2-3 years (Engers and Gabbert, 2000; Kennecke *et al*., 2010; Shibue and Weinberg, 2011; Qiao *et al*., 2022). Bone metastasis falls into three different categories, namely osteolytic (involves bone resorption), osteoblastic (involves deposition of new bone), and mixed (involves both formation and resorption of bone), with the vast majority of breast cancer-associated metastasis being osteolytic in nature (>70%) (Chen, Sosnoski, and Mastro, 2010). Metastasis to the bone is complex process that involves chemotactic migration and attachment of circulating tumor cells to the bone surface, and subsequent interactions between cancer cells and various stromal cells in the bone microenvironment that lead to a vicious cycle of bone remodeling. Under normal conditions, bone-forming osteoblasts (OBs) produce macrophage colony-stimulating factor (MCSF) and receptor activator for nuclear factor κB (NFκB) ligand (RANKL) which stimulate proliferation and differentiation of myeloid progenitor cells from the bone marrow into macrophage/monocyte lineage cells and their subsequent differentiation/maturation into osteoclasts (OCLs), a type of cells that is responsible for degradation of mineralized bone. However, OBs also produce osteoprotegerin (OPG) which binds to and sequesters RANKL thereby limiting OCL activity and regulating the levels of bone resorption (Tsuda *et al*., 1997; Yasuda *et al*., 1998; Boyce and Xing, 2008; Weidle *et al*., 2016). The balancing act of OBs and OCLs is dysregulated in the presence of tumor cells where osteoclastogenesis and OCL survival are greatly enhanced by increased secreted levels of RANKL and other soluble factors (e.g. IL6, IL11, IL1, TNF, PTHrP) contributed by tumor cells, osteocytes, OBs and immune cells (Kang *et al*., 2003; Nguyen, Bos, and Massagué, 2009; David Roodman and Silbermann, 2015). Increased bone resorption promotes the release and activation of ECM-bound growth factors and cytokines, further amplifying this loop and stimulating tumor cell proliferation and metastatic colonization of tumor cells (Casimiro *et al*., 2016).

Myocardin-related transcription factors (MRTFs) are key coactivators of serum-response factor (SRF) that link actin dynamics to SRF-mediated transcription of a wide variety of genes including SRF itself and many others which play important roles cytoskeleton remodeling, cell adhesion and ECM regulation (Olson and Nordheim, 2010; Gasparics and Sebe, 2018; Gau and Roy, 2018). There are two major isoforms of MRTF, namely MRTF-A (also known as MKL1) and MRTF-B (also known as MKL2). Both MRTF isoforms are ubiquitously expressed, share structural and functional similarities, and compensate for each other in most loss-of-function settings. However, phenotypic differences between mice deficient in MRTF-A vs-B suggest that these isoforms are not completely functionally redundant (Oh, Richardson, and Olson, 2005; Sun *et al*., 2006).

Consistent with the role of the MRTF/SRF transcriptional axis in regulating the expression of many structural and regulatory components of actin cytoskeleton, depletion of either MRTFs or SRF causes defects in actin cytoskeleton and cell migration of both normal and cancer cells (including breast cancer cells) *in vitro* and *in vivo* (comprehensively reviewed in (Gau and Roy, 2018)). We recently showed that nuclear localization of MRTF positively correlates with the malignant traits of tumor cells in human breast cancer, and further linked actin-binding protein mammalian Diaphanous 2 (mDia2) as a mediator of MRTF-A dependent regulation of breast cancer cell migration (Eder *et al*., 2024). Several studies have underscored the importance of MRTF-SRF function for metastatic colonization of tumor cells. For example, the Treisman group showed that silencing the expression of either MRTF or SRF dramatically impairs experimental lung colonization ability of breast cancer and melanoma cells *in vivo* (Medjkane *et al*., 2009). Supporting these findings, we recently demonstrated that MRTF-depletion induces an outgrowth-arrest phenotype in breast cancer cells, and conversely, overexpression of either MRTF-A or MRTF-B accelerates initiation and progression of breast cancer cells on soft ECM substrate. Our *in vivo* studies further showed that SRF’s interaction with MRTF is critical for the intrinsic tumor-initiating ability and post-extravasation metastatic outgrowth of breast cancer cells in immunodeficient mice (Gau *et al*., 2022). Interestingly, when MRTF is overexpressed in breast cancer cells, increased actin polymerization and cell stiffening also increases susceptibility of tumor cells to cytotoxic T cell attack thereby limiting lung colonization of tumor cells through immune surveillance (Tello-Lafoz *et al*., 2021). Collectively, these findings suggest that a balanced MRTF activity in tumor cells is perhaps most efficient for their metastatic colonization ability in the presence of a functional immune system.

MRTF’s role in metastatic colonization of cancer cells has been primarily investigated in the context of soft organs to date, and those studies focused exclusively on tumor-intrinsic impact of perturbation of MRTF. Bone represents a unique structural organ in terms of its physiochemical characteristics (ECM composition and stiffness) and parenchymal/non-parenchymal cellular compositions. The objective of the present study was to investigate tumor-extrinsic impact of perturbation of MRTF, specifically examining its potential role in tumor-cell directed paracrine modulation of OCL differentiation.

## RESULTS

### MRTF plays a critical role in tumor cell-directed modulation of osteoclast differentiation

To investigate whether MRTF has any role in tumor-cell directed paracrine modulation of OCL differentiation, we conducted studies with two metastatic triple-negative breast cancer [TNBC] cell lines (TNBC – lacks the expression of estrogen receptor (ER), progesterone receptor (PR) and human epidermal growth factor receptor (HER)) namely, MDA-MB-231 (MDA-231– a human TNBC cell line) and D2.A1 (a murine TNBC cell line). Since loss-of-expression of one isoform of MRTF can be functionally compensated by the other isoform, we genetically engineered GFP/luciferase-expressing sublines of MDA-231 and D2.A1 cells for stable co-suppression of MRTF isoforms using a single shRNA that targets an mRNA region common to both isoforms. Corresponding non-targeting control shRNA-transfectants served as controls. Depending on the cell line and the isoform of MRTF, we achieved an average of 60-90% reduction of MRTF isoforms at the protein level in the MRTF-A/B shRNA (referred to as MRTF-knockdown hereon) sublines vs their respective controls (**Figs 1A-B**). Reduction of the total MRTF level diminished the rate of proliferation of both breast cancer cell lines in 2D culture, although the effect was more prominent in D2.A1 cells likely due to better knockdown efficiency in this cell line (**Fig 1C**). For OCL differentiation studies, we first differentiated bone marrow progenitor cells harvested from female FVB mice into macrophages (denoted as bone marrow-derived macrophages or BMDMs in short) by MCSF (macrophage colony stimulating factor) treatment. Subsequently, we differentiated BMDMs into OCLs by co-treatment of RANKL and MCSF with or without supplementation of conditioned media (CM) harvested from 2D cultures of control vs stable MRTF-knockdown breast cancer (MDA-231 and D2A1) cells (the experimental schematic is shown in **Fig 2A**). BMDMs without RANKL exposure served as the negative control group in these experiments. Although stable knockdown of MRTF reduced proliferation of both breast cancer cell lines in 2D culture, the total protein concentration in the CM was not significantly affected by MRTF depletion in either of the two cell lines (**supplementary Fig S1**). Induction of expression of Tartrate-resistant alkaline phosphatase (TRAP - encoded by the *ACP5* gene), an enzyme involved in hydrolysis of bone matrix, is a hallmark of OCL differentiation, and the phosphatase activity of TRAP (a proxy marker of osteoclastic bone resorption (Kirstein, Chambers, and Fuller, 2006)) can be visualized in cells by a well-established colorimetric assay (Ethiraj *et al*., 2022). Therefore, we performed end-point colorimetric TRAP activity staining of cultures to identify and score the relative % of TRAP-negative (undifferentiated) vs TRAP-positive cells with either 1-2 nuclei (denoted as OCL-precursor or OCLp in short) or multi-nucleated (presence of 3 or more nuclei – characterized as mature OCL) phenotypes. Representative images of TRAP-stained cells from these experiments are shown in **Fig 2B**. In general, we observed a trend of reduced total cell counts of BMDM cultures when they were exposed to tumor cell-derived CM; however, MRTF-depletion in tumor cells per se did not have any additional effect (**supplementary Fig S2**). Because of differences in the total cell counts of BMDM cultures between the groups, to compare the intrinsic differentiation potential of BMDMs for the different treatment groups we quantified the relative % of TRAP-negative (undifferentiated), TRAP-positive OCLp, and TRAP-positive mature OCL in each group (summarized in **Fig 2C**), and normalized % of mature OCLs in a given group to the same calculated for RANKL-alone treatment group, which we termed as “OCL differentiation index” (graphically displayed in **Fig 2D**). In both cell line experiments, RANKL addition dramatically promoted OCL differentiation of BMDMs as expected. RANKL-induced OCL differentiation index was further enhanced when BMDMs were exposed to the CM harvested from control shRNA cells, and this trend was not only reversed but was also actively repressed when MRTF expression was stably silenced in breast cancer cells (**Fig 2D**). When we compared the relative proportion of TRAP-negative vs TRAP-positive (i.e. the combined pool of OCLp and OCL) cells between various RANKL-stimulation groups, % TRAP-negative cells in BMDM culture exposed to CM of MRTF-depleted cells (∼43-47%) was strikingly higher than other two RANKL-stimulated groups (∼1-4%) (**Fig 2C**). These data suggest that the secreted content from MRTF-depleted cells dramatically prevents the early aspect of OCL differentiation. Furthermore, although the total % of TRAP-positive cells (∼95-98%) of the RANKL-stimulated group was not enhanced by the addition of CM derived from control shRNA-expressing tumor cells, we noted an increasing trend in the ratio of OCL-to-OCLp pool (**Fig 2C**). These findings could further suggest that tumor-cell derived factors somehow accelerate the OCL maturation process.

**Figure 1.**
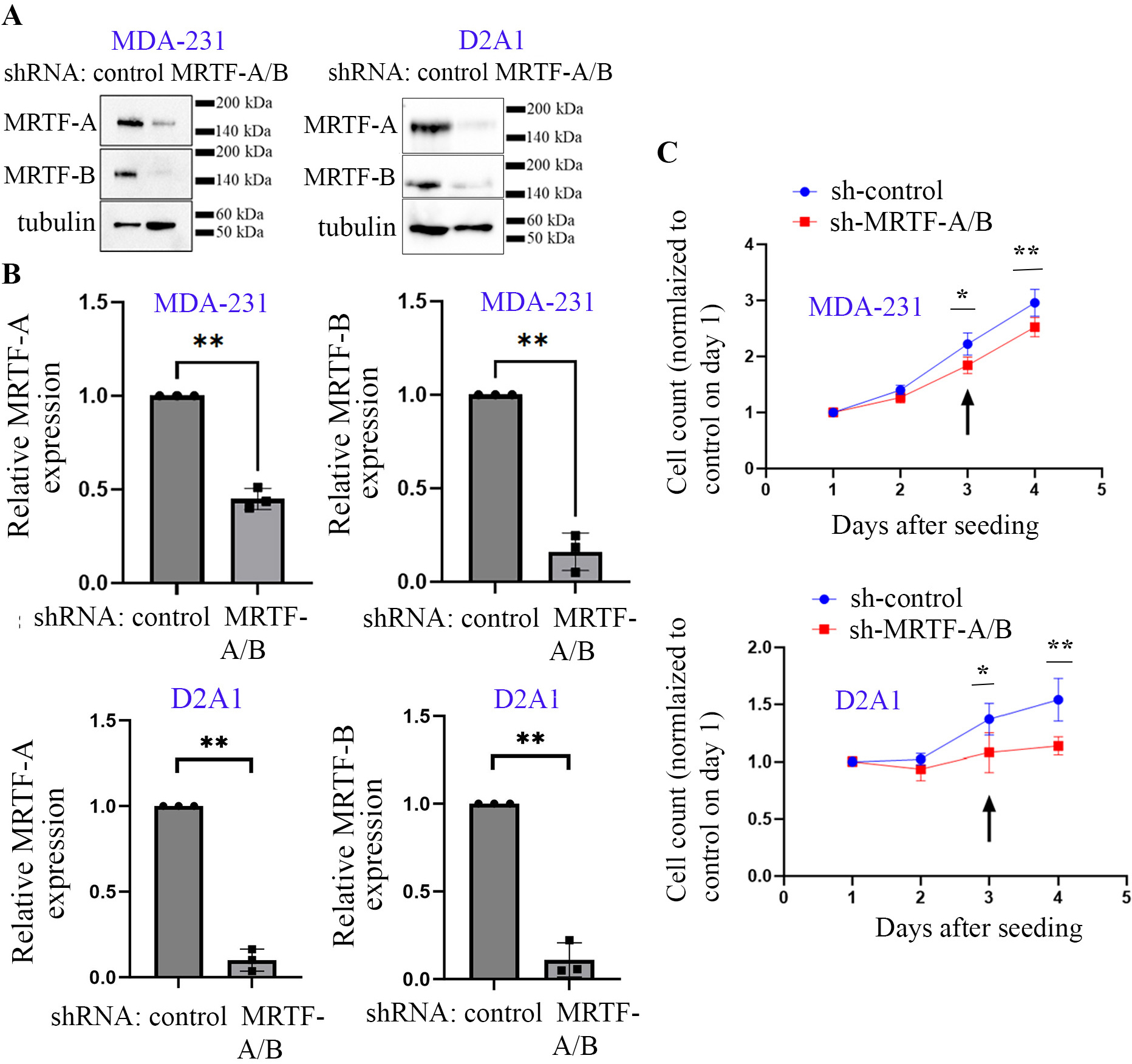
Dual knockdown of MRTF expression in breast cancer cell lines. **A-B)** Representative immunoblots of total cell lysates of MDA-231 and D2.A1 cells (*panel A;* tubulin blot: loading control) and quantifications of immunoblot data (*panel B*) demonstrating shRNA-mediated stable knockdown of MRTF isoforms in both cell lines. **C)** Proliferation curves of control vs stable MRTF-A/B knockdown MDA-231 and D2A1 cells in 2D tissue culture (relative to the value measured for control groups of cells on day 1); arrow indicates the day conditioned media (CM) was harvested for subsequent OCL differentiation experiments. All data are summarized from 3 independent experiments; *: p<0.05; **: p<0.01).

**Figure 2.**
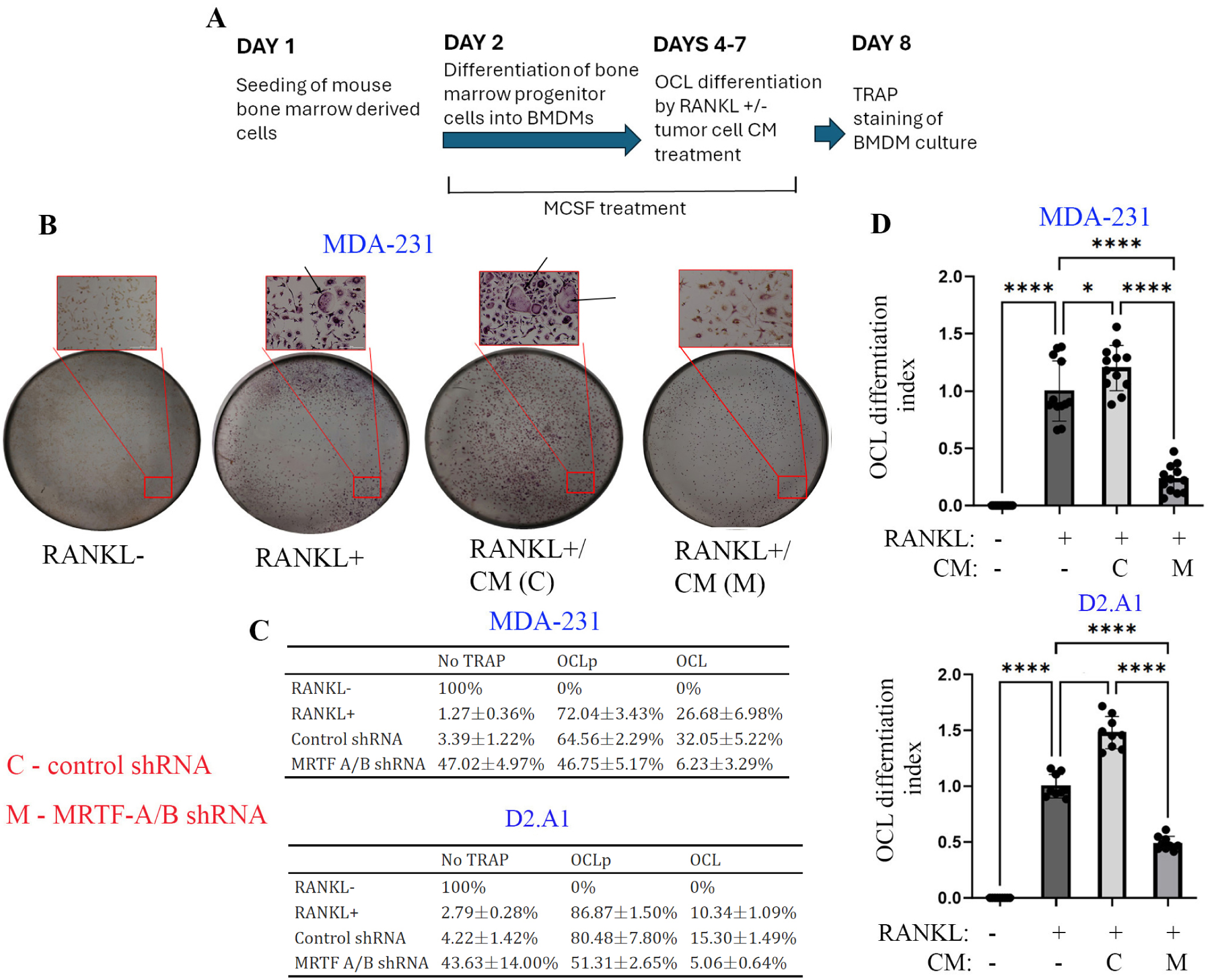
Tumor cell-derived factors promote RANKL-induced OCL differentiation of BMDMs in an MRTF-dependent manner. **A)** Schematic of the experimental protocol of RANKL-induced OCL differentiation with or without tumor cell CM supplementation. **B)** Representative images of TRAP-stained BMDM cultures in the absence or presence of RANKL (with or without MDA-231 CM supplementation; C-control shRNA; M-MRTF-A/B shRNA; scale bar in the inset images – 200 μm). Large multinucleated cells in the inset images represent mature OCLs. **C)** Summary of relative % of TRAP-negative vs TRAP-positive (1-2 nuclei) vs TRAP-positive (3 or more nuclei) cells in the BMDM cultures for the indicated treatment groups. **D)** Quantifications of OCL differentiation indices of BMDM cultures for the indicated treatment groups (data summarized from 4 and 3 independent experiments for CM treatment from MDA-231 and D2.A1 cells respectively (* p<0.05, **: p<0.01, and **** p<0.0001).

DC-Stamp (dendritic cell-specific transmembrane protein) and Cathepsin-K (a protease) are major regulators of osteoclastogenesis, and deficiency of either is linked to decreased differentiation and/or activity of OCLs (Saftig *et al*., 2000; Yagi *et al*., 2005). Specifically, DC-stamp is responsible for cell-cell fusion giving rise to the multinucleated mature OCL phenotype (Yagi *et al*., 2005). Cathepsin-K plays a key role in proteolytic cleavage and conversion of the inactive TRAP precursor protein (the initially synthesized form) into an active form (Ljusberg *et al*., 2005). Consistent with our OCL differentiation readout data, quantitative RT-PCR analyses showed that RANKL-induced expressions of both DC-stamp and Cathepsin-K in BMDMs are further enhanced by MDA-231 secreted factors in an MRTF-dependent manner (**Supplementary Fig S3**).

To further establish the specificity of these knockdown findings, we undertook two different approaches. First, we repeated these OCL differentiation studies in a transient knockdown setting of MRTF isoforms in MDA-231 cells. For these experiments, we combined individual pooled MRTF-A-specific and MRTF-B-specific siRNAs targeting their coding regions that are different from the common region targeted by the shRNA in our stable knockdown experiments. Although exposure to CM harvested from siRNA-transfected tumor cells did not reduce the proliferation of BMDM cultures as we observed in the case of stable knockdown setting, the overall trend of tumor cell-derived factors promoting RANKL-induced OCL differentiation in an MRTF-dependent manner was also recapitulated in the transient knockdown setting (**supplementary Fig S4**). Second, as a complementary approach, we conducted OCL differentiation studies in an overexpression setting involving either wild-type (WT) or select functional mutant forms of MRTF-A. Although SRF-activation is the most well-studied function MRTF, both MRTF isoforms also contain a SAP (SAF-A/B, Acinus and PIAS – named after the proteins the domain was originally discovered in) domain commonly found in other DNA-binding proteins. At least in a transient transfection setting, deletion of the SAP domain does not affect MRTF’s ability to activate SRF (Cen *et al*., 2003). However, overexpression of the SAP-domain-deficient mutant form of MRTF-A induces transcriptomic changes in cells that are distinct from those elicited by overexpression of K237A/Y238A/H239A/Y241A-MRTF-A (an SRF-binding deficient mutant of MRTF-A) which led to a postulate that MRTF may also have gene-regulatory function utilizing its SAP domain in an SRF-independent manner (Asparuhova *et al*., 2011; Gurbuz *et al*., 2014; Asparuhova *et al*., 2015). We previously engineered GFP/luciferase expressing sublines of MDA-231 cells for doxycycline (dox)-inducible overexpression of either WT-MRTF-A or K237A/Y238A/H239A/Y241A-MRTF-A (denoted as ΔSRF-MRTF-A) or SAP-domain-deleted mutant of MRTF-A (denoted as ΔSAP-MRTF-A). Utilizing these cell lines, we showed that MRTF-A overexpression-induced stimulation of various tumor cell activities (e.g. migration, invasion, single-cell outgrowth, tumor-initiation, metastatic outgrowth, and colonization) are inhibited dramatically particularly by mutations that disrupt SRF’s interaction of MRTF-A, mimicking previously reported phenotypes in knockdown settings (note that invasion was also negatively affected by SAP-domain deletion) (Gau *et al*., 2022; Eder *et al*., 2024). Therefore, at least the ΔSRF-MRTF-A mutant cell line appears to be as a reasonable loss-of-function model in an overexpression setting, prompting us to further examine whether CM derived from these various MRTF-A overexpressing sublines has any differential effect on RANKL-induced OCL differentiation. As demonstrated in **supplementary Fig S5**, overexpression of WT-MRTF-A did not further potentiate OCL differentiation when compared to control; however, CM derived from either ΔSRF- or ΔSAP-MRTF-A overexpressing sublines actively repressed OCL differentiation mimicking our knockdown experimental results. Note that the total cell count of BMDM culture was not significantly different between experimental conditions that involved treatment of CM derived from any of the MRTF-A overexpression cultures. Collectively, these data demonstrate that MRTF activity in tumor cells plays a key role in promoting OCL differentiation and that both SAP-domain- and SRF-related activities of MRTF are important for tumor-cell directed paracrine regulation of OCL differentiation.

### MRTF depletion in breast cancer cells downregulates multiple pro-osteoclastogenic cell secreted factors

To gain insight into how loss of MRTF expression in breast cancer cells dampens RANKL-induced OCL differentiation in a paracrine manner, we first compiled a signature of 21 genes based on two previously published findings (Kang *et al*., 2003; Savci-Heijink *et al*., 2015) that are associated with bone metastasis in human breast cancer (**supplementary Table S1**). We then queried this gene set against previously published differentially expressed genes (DEGs) of 2D cultures of MDA-231 cells upon stable co-silencing of MRTF isoforms (Medjkane *et al*., 2009). Based on this query, we identified three bone metastasis-associated genes that are downregulated in mRNA expression upon MRTF silencing, including CTGF/CCN2 (consistent with a previous finding in endothelial cells in an overexpression setting (Hinkel *et al*., 2014)), IL11, and FGF5. From our recently published RNA-seq analyses of 2D and 3D cultures of GFP/Luc expressing sublines of MDA-231 cells with or without overexpression of WT-vs functional mutants of MRTF-A (GSE253047) (Eder *et al*., 2024), we were able to further verify that expressions of all three genes are increased in MDA-231 cells when WT MRTF-A is overexpressed (**Fig 3A**) although the effect of functional disruption of MRTF on its ability to induce these genes was found to be gene-specific. While MRTF-A’s ability to induce CTGF expression at the transcript level requires both SRF- and SAP-domain-related functions, induction of IL11 and FGF5 were selectively dependent on SRF- and SAP-domain-related functions of MRTF-A, respectively. In general, there was good concordance in the patterns of expression of these three genes between the 2D and the 3D culture conditions. Interestingly, all three genes (CTGF, IL11 and FGF5) encode for cell-secreted factors that are known to promote OCL differentiation and/or activity (McCoy *et al*., 2013; Aukes *et al*., 2017; Choi *et al*., 2020). However, when we analyzed three large human breast cancer transcriptome databases (METABRIC [n=1981] (Pereira *et al*., 2016), TCGA [n=1208] (Ciriello *et al*., 2015), and SCANB [n=3207] (Brueffer *et al*., 2020); n represents the number of clinical samples), only CTGF expression showed a strong positive correlation (spearman’s correlation coefficient ∼0.6) with the MRTF-A gene signature score (constructed based on the gene signature variation analyses (GSVA) as detailed in our recent report (Hänzelmann, Castelo, and Guinney, 2013; Eder *et al*., 2024)) in primary tumors consistently in all three transcriptome databases (**Fig 3B**). Furthermore, when we queried the TCGA database, only MRTF-A (and not MRTF-B) has a significant positive association with CTGF expression at the mRNA level in human breast cancer (**Fig 3C**). Based on these clinical correlation findings, we chose to further interrogate CTGF as a candidate mediator of MRTF-dependent paracrine regulation of OCL differentiation.

**Figure 3.**
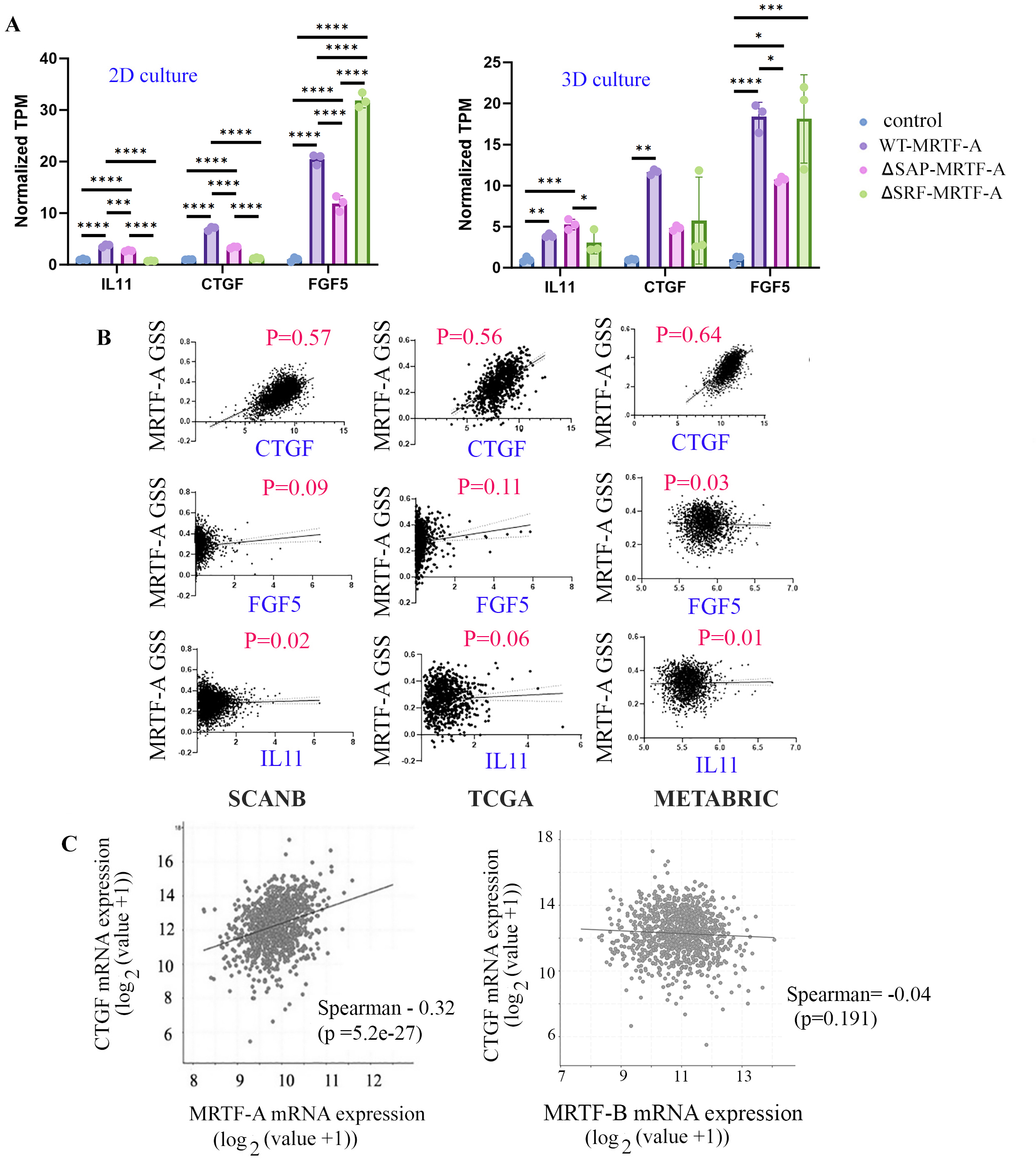
Identification of candidate bone metastasis-associated genes inducible by MRTF. **A)** TPM (transcripts per million – normalized to housekeeping genes) of candidate bone metastasis associated genes (IL11, CTGF, FGF5) based on RNA-sequencing data of 2D and 3D cultures of MDA-231 cells subjected to acute doxycycline-induced overexpression of wild-type (WT) vs various mutant forms of MRTF-A. **B)** Correlation between select genes and MRTF-A gene signature score (GSS) in TCGA, MRTABRIC and SCANB databases (Spearman’s co-efficient (P) are included for each of these analyses). **C)** Correlation between CTGF and MRTF-A/B mRNA expression in human breast cancer samples (based on TCGA data analyses – data downloaded from CBioportal platform; * p<0.05, **: p<0.01, and **** p<0.0001).

Although overexpression studies demonstrate MRTF-A dependent stimulation of CTGF expression (Hinkel *et al*., 2014), given that CTGF expression can be also regulated by other transcription factors (e.g. YAP (Cheng *et al*., 2018)), to what extent MRTF is required for maintaining the basal CTGF expression at the protein level is not known. We found that MRTF knockdown in both MDA-231 and D2A1 cells results in a dramatic reduction (∼80-90% depending on the cell line) in CTGF expression at the protein level, suggesting that MRTF is essential for maintaining the basal expression of CTGF (**Fig 4A-B**). We also compared the CTGF expression between the various MRTF-A overexpressing sublines of MDA-231 cells relative to the control subline, the results of which are shown in **Fig 4C**. Consistent with the knockdown experimental results, overexpression of WT-MRTF-A increased CTGF expression relative to control in all three independent experiments with the p-value (=0.07) close to statistical significance (note that the variation in the fold-change accounts for the p-value not reaching statistical significance). MRTF-A-induced increase in CTGF expression was completely reversed upon functional disruption of either SRF’s interaction or SAP-domain function. CTGF expression in ΔSRF-MRTF-A expressers was the least among all four groups indicating a critical importance of SRF’s interaction of MRTF is maintaining the basal cellular CTGF expression.

**Figure 4.**
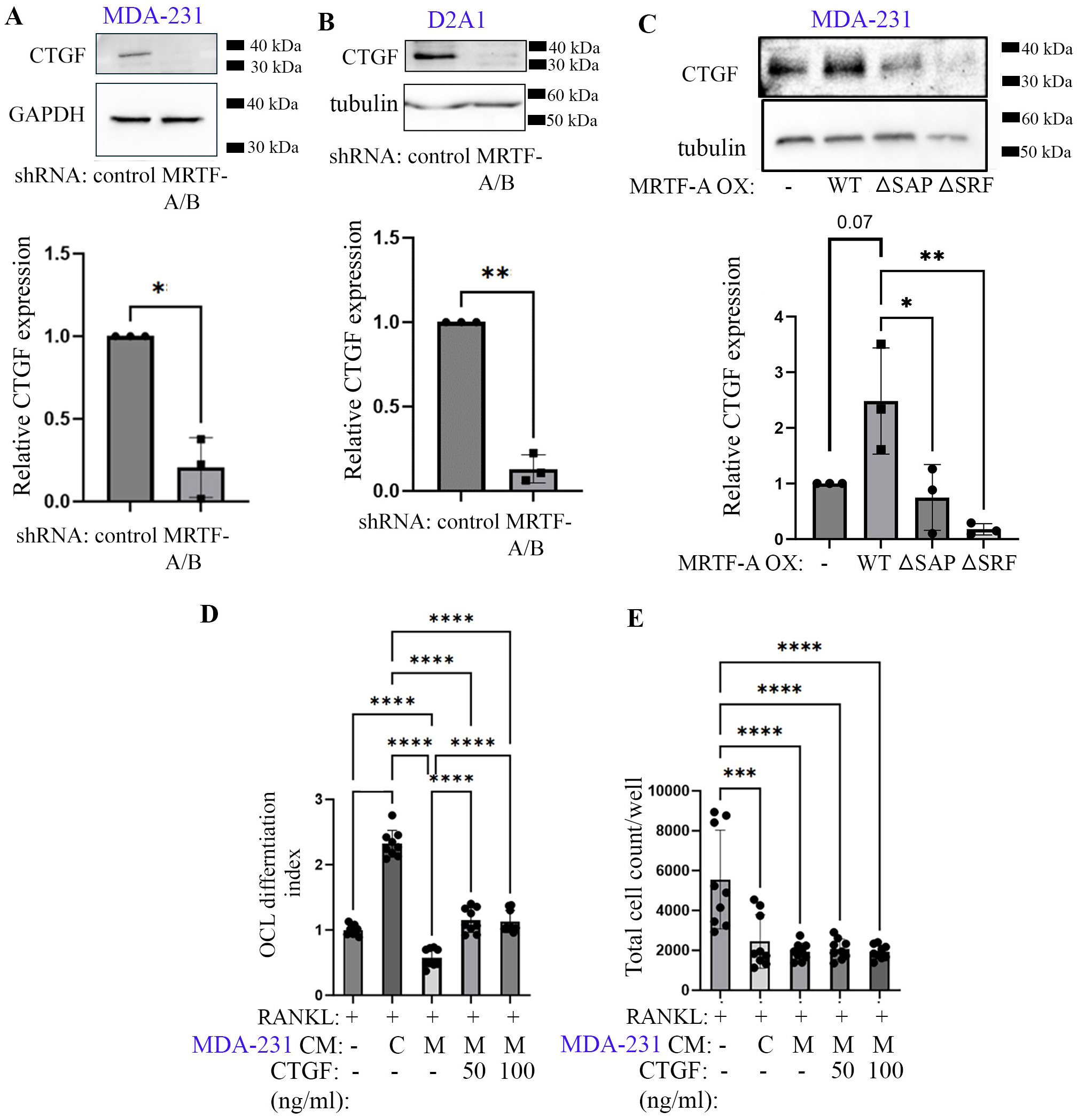
CTGF is a mediator of MRTF-dependent paracrine regulation of OCL differentiation by breast cancer cells. **A-B)** CTGF immunoblots and the associated quantification of the immunoblot data of total cell lysates of MDA-231 (*panel A*) and D2A1 (*panel B*) cells with or without stable knockdown of MRTF isoforms (GAPDH and tubulin blots serve as loading controls). **C)** Representative CTGF immunoblot and the associated quantification control vs various MRTF-A (WT and functional mutants) overexpressing sublines MDA-231 cells. **D-E)** Quantification of OCL differentiation index (*panel D)* and total cell count (*panel E*) of RANKL-stimulated BMDM cultures without or with supplementation of MDA-231 CM and/or recombinant CTGF at the indicated concentrations. All data presented here are summarized from 3 independent experiments; *: p<0.05, **: p<0.01, *** p<0.001, and **** p<0.0001).

Next, we asked whether CTGF supplementation can rescue OCL differentiation defect of BMDMs in the setting of MRTF knockdown setting. Previous studies reported CTGF supplementation at 50-100 ng/ml boosts RANKL-induced OCL differentiation in BMDMs (derived from IJR strain male mouse) and RAW264.7 (a Balb/c mouse-derived tumorigenic macrophage cell line) cells (Nishida *et al*., 2011; Choi *et al*., 2020). In a rescue setting, CTGF addition at 50 ng/ml concentration reversed the dampening of RANKL-induced OCL differentiation of BMDMs induced by CM of MRTF-silenced MDA-231 cells (note that the total cell count in the BMDM culture was not impacted by CTGF addition) (**Figs 4D-E)**. Increasing CTGF concentration from 50 to 100 ng/ml did not result in any further increase in OCL differentiation in the BMDM culture. These data suggest that CTGF is an important paracrine mediator of pro-osteoclastogenic action of MRTF in breast cancer cells.

Since CTGF supplementation was not sufficient to abrogate the difference in the OCL differentiation index of BMDMs between the two (i.e. control vs MRTF-knockdown) CM exposure settings (**Figs 4D-E**), we postulated that MRTF depletion in breast cancer cells may alter secretion of additional pro- and/or anti-osteoclastogenic factors. Many immunomodulatory cytokines and chemokines play a prominent role in both positively and negatively regulating proliferation, survival and differentiation of OCL (Amarasekara *et al*., 2018b). Therefore, we performed Luminex analyses of CM harvested from control vs MRTF knockdown cultures of MDA-231 cells and D2A1 cells to profile a panel of 48 human and 32 mouse cytokines/chemokines, respectively (**supplementary Table S2)** lists the cytokines/chemokines probed by each of these multiplexed Luminex panels with the common ones indicated by red). We detected a total of 13 differentially abundant factors (within statistical significance) between the two groups with 7 factors upregulated (IL1RA, IP10/CXCL10, IL1β, TGFα, fractalkine/CX3CL1, IL27, RANTES/CCL5) and 6 factors downregulated (IL8, G-CSF, FGF2, IL3, IL17E/IL25, FLT3L) in the CM of MDA-231 cells upon MRTF knockdown within statistical significance (**Fig 5A**). Among these factors, upregulation of anti-osteoclastogenic factors (IL1RA and IL27) and conversely, downregulation of pro-osteoclastogenic factors (IL8, G-CSF, and FLT3L) are consistent with the repression of OCL differentiation in an MRTF knockdown setting. Similarly, analyses of D2.A1 CM revealed a total of 10 differentially abundant factors between the two groups with 1 factor upregulated (IL7) and 9 factors downregulated (MIP1a/CCL3, LIX/CXCL5, GM-CSF, MCP1/CCL2, Eotaxin, IL1α, IL6, G-CSF) upon MRTF knockdown within statistical significance (**Fig 5B**). Among these factors, reduction in G-CSF, CCL2, CCL3, IL1α, IL6 and Eotaxin are directionally consistent with repression of OCL differentiation in the MRTF knockdown setting. Collectively, these data demonstrate that loss of MRTF expression in breast cancer cells has a broad impact on the secretion of OCL-regulatory factors. The general lack of consensus in differentially abundant cytokines/chemokines between these two sets of Luminex data was not surprising because of inherent differences between human vs mouse breast cancer cell lines. However, reduced secreted level of G-CSF and potential downregulation of IL1 signaling (as suggested by reduction of secreted IL1 or upregulation of secreted IL1 receptor antagonist [IL1RA]) upon MRTF depletion seemed to be a common feature in both breast cancer cell lines. It is worth noting that we did not find evidence for changes in the mRNA levels of any of these immunoregulatory factors when we queried previously published microarray data (Medjkane *et al*., 2009) and our own RNA-seq data of MDA-231 cells in knockdown and overexpression settings of MRTF, respectively. This suggests that MRTF-dependent changes in these immunomodulatory factors are most likely not through direct transcriptional control.

**Figure 5.**
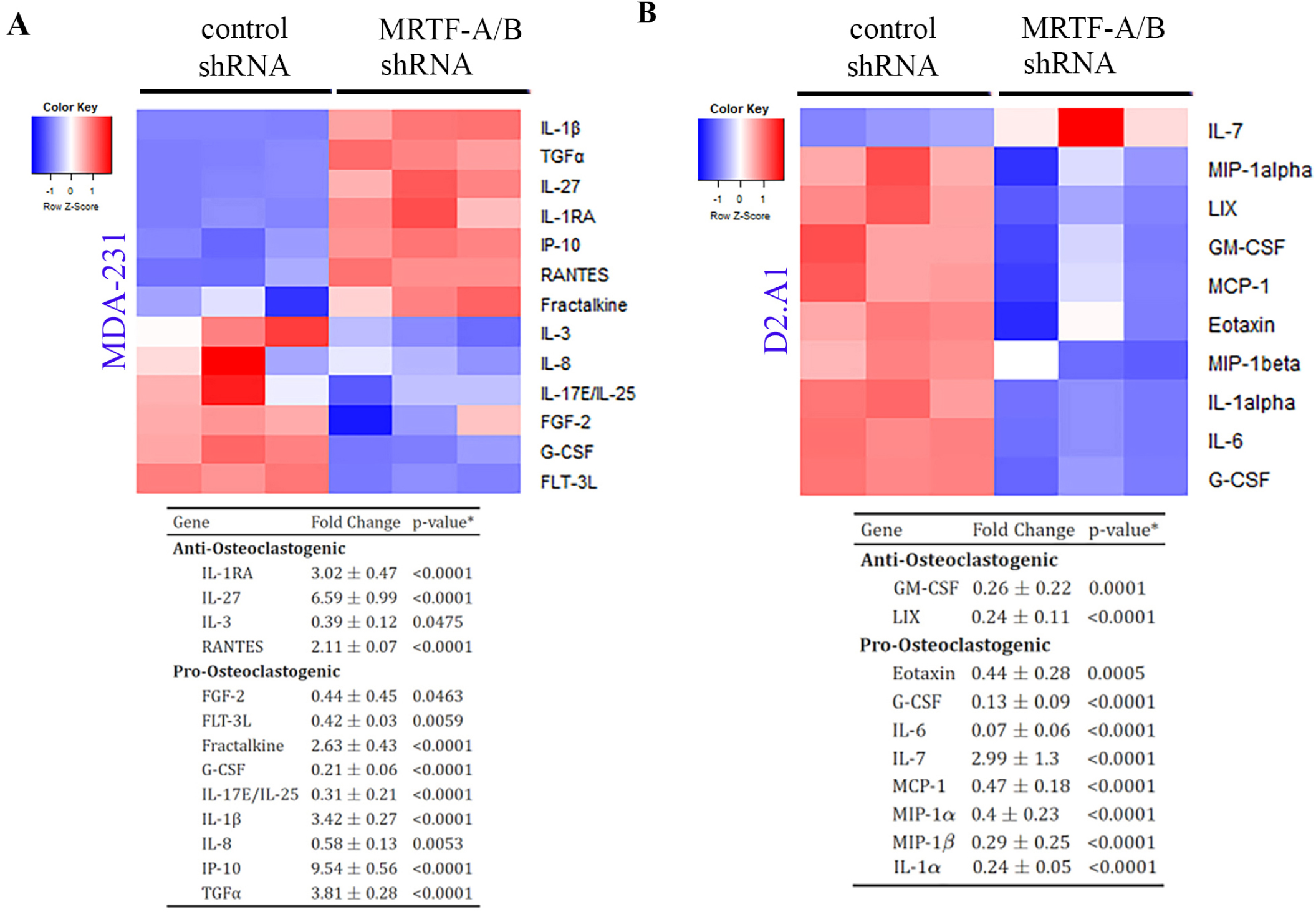
MRTF-dependent changes in secreted immunomodulatory factors. Heat plots of differentially abundant cytokines/chemokines in the CM of control vs stable MRTF knockdown (KD) MDA-231 (*panel A*) and D2A1 (*panel B*) cultures. Fold changes and p-values for differentially abundant cytokines/chemokines with pro-vs anti-osteoclastogenic activities are indicated below.

### MRTF depletion in breast cancer cells diminishes OCL abundance and experimental bone colonization in vivo

To query MRTF’s role in bone colonization of breast cancer cells, we first performed intracardiac injection-based experimental metastasis assays with control *vs* MRTF knockdown MDA-231 cells in immunodeficient NOD-SCID-gamma (NSG) mice. Although orthotopic models represent the entire cascade of events of metastasis, we chose experimental metastasis assays to exclusively assess the impact of loss of MRTF expression on metastatic growth of breast cancer cells in bone without confounding effects of MRTF-dependent changes in primary tumor growth and/or tumor cell dissemination. Intracardiac experimental metastasis assay allows for widespread metastasis of cancer cells including to the bone. Use of NSG mice also enabled us to decouple the secondary effect of MRTF on immune surveillance in metastatic colonization process. (Gau *et al*., 2022; Eder *et al*., 2024)Bioluminescence imaging (BLI) revealed that 6 out 8 animals (75%) animals inoculated with control MDA-231 subline exhibited skeletal metastasis in the long bones within 11 days after injection, compared to only 1 out 8 (12.5%) animals injected with MRTF knockdown cells presenting the same (**Fig 6A**; BLI quantification and skeletal metastasis confirmation by bone histology are shown in **6B-C**). Chemotactic migration, adhesion, and proliferation are all essential for efficient bone colonization of tumor cells, and actin cytoskeleton plays important roles in all these processes. We previously showed that overexpression of MRTF-A stimulates actin polymerization, random migration and invasion of MDA-231 cells (Gau *et al*., 2022; Eder *et al*., 2024). Consistent with these findings, MRTF knockdown diminished the overall F-actin content in MDA-231 cells (**supplementary Fig S6**). Transwell migration experiments revealed that MRTF knockdown and overexpression inhibited and stimulated serum-induced chemotactic migration of MDA-231 cells (**supplementary Fig S7**). Although MRTF knockdown did not impair adhesion of MDA-231 cells on calcium-phosphate (CaP)-coated tissue-culture substrate (mimics mineralized bone matrix), it dramatically inhibited outgrowth of MDA-231 cells when seeded on devitalized bone surface prepared from mouse calvaria (**supplementary Fig S8**). Since we also do not see any evidence of cell death in MRTF-silenced cells in the cell culture, these data further underscore a major proliferation defect of breast cancer cells induced by the loss of MRTF expression.

**Figure 6.**
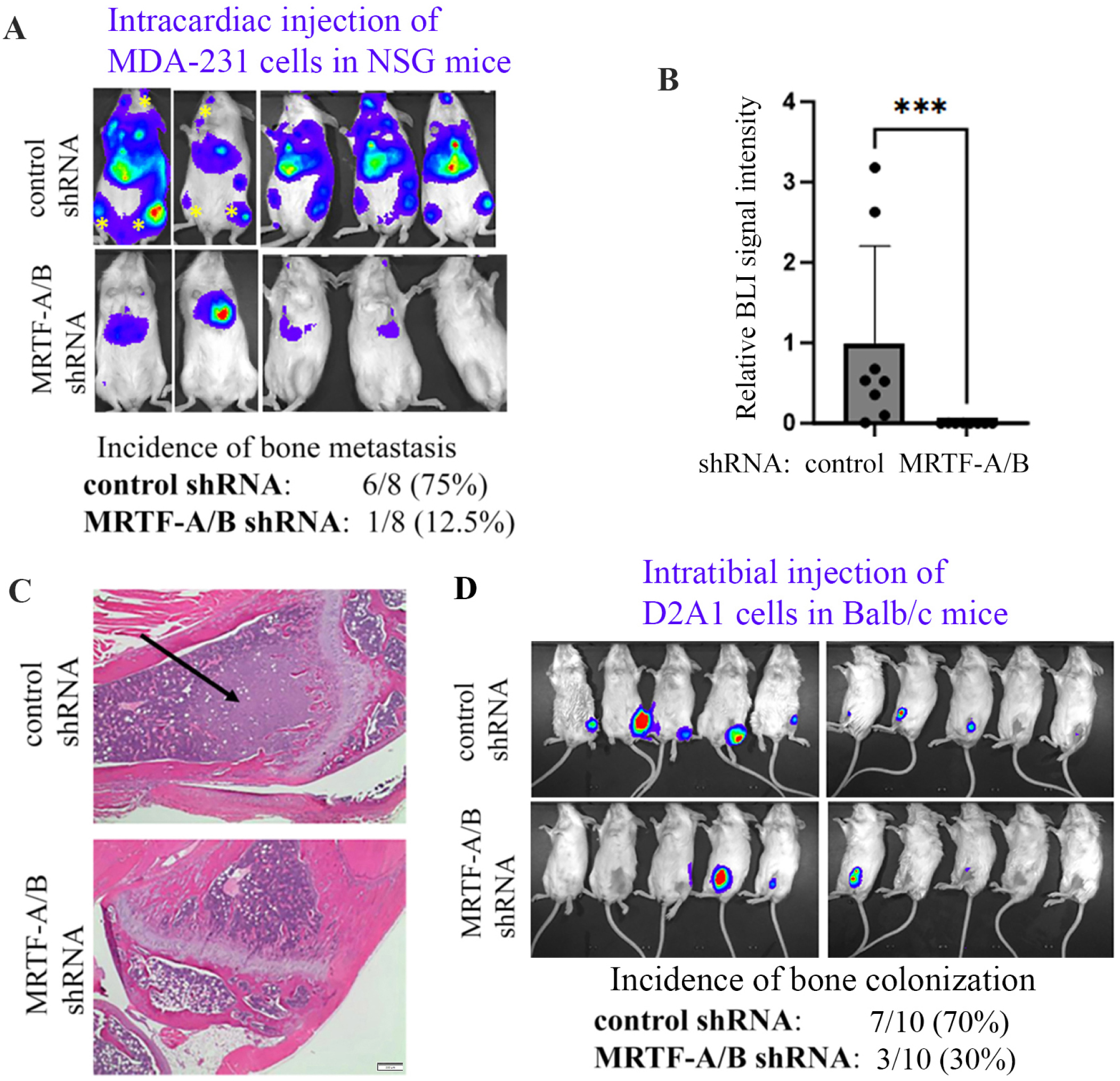
MRTF depletion impairs experimental bone colonization of breast cancer cells *in vivo*. **A-C)** Representative endpoint BLI images (*panel A*) NSG mice 2 weeks after intracardiac injection of control vs MRTF-A/B knockdown MDA-231 cells (asterisks indicate skeletal metastasis). Panel B summarizes the distant metastasis quantification based on the BLI signaling readouts (data normalized to the control group). Hematoxylin and Eosin staining of BLI signal-positive bones in panel C confirms presence of metastases (indicated by the arrow) harvested from mice injected with control shRNA-bearing cells. Scale bar represents 200 µm. **D)** BLI images of immunocompetent Balb/c mice 1 week after intratibial injection of control vs MRTF-A/B knockdown D2.A1 cells.

Although the intracardiac injection method is widely used for assessing experimental metastatic colonization of cancer cells, the overall metastatic burden in this *in vivo* assay is also influenced by tumor cell survival in the circulation as well as extravasation ability of tumor cells at the metastatic site. Furthermore, while *in vivo* studies involving human cancer cells necessitate the use of immunodeficient animals, since immune cells have a multitude of effects on OBs and OCLs in bone metastatic microenvironment thereby dictating the outgrowth behavior of cancer cells (Chen *et al*., 2024), fully immunocompetent mouse models are more physiologically relevant model systems. Therefore, in our next set of experiments, we performed direct intratibial injection of control vs MRTF-knockdown D2A1 cells in syngeneic immunocompetent Balb/c mice. Consistent with our findings of MDA-231 study, 70% (7 out of 10) of mice subjected to intratibial injection of control D2A1 cells presented with metastatic growth in the bone (detectable by BLI signal) within 7 days after injection, compared to only 30% (3 out 10) mice injected with MRTF knockdown cells (**Fig 6D**). Furthermore, corroborating our in vitro findings, TRAP-staining of a subset of bones from these experiments also revealed prominently lower OCL abundance in MRTF-knockdown-vs control D2A-injected mouse tibia (**Fig 7**). Collectively, these data underscore MRTF’s importance in OCL differentiation and bone colonization of breast cancer cells.

**Figure 7.**
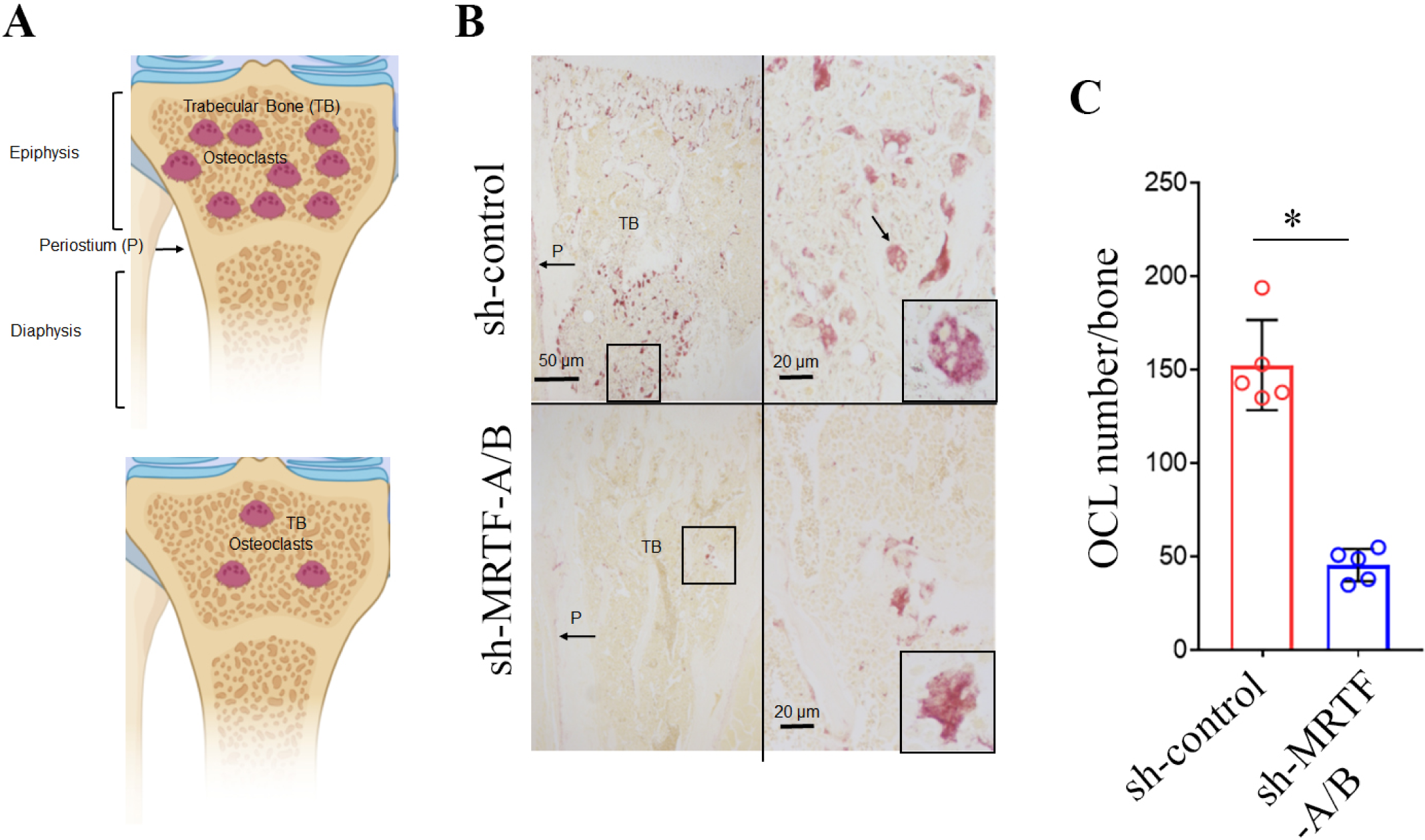
MRTF’s presence in tumor cells impacts OCL abundance in vivo. **A)** Diagram illustrating the anatomy of mouse tibias showing the area where an increased number of OCLs was found (created with BioRender). **B)** Upper left; histological image of a mouse tibia injected with control shRNA-expressing D2.A1 breast cancer cells showing a high number of OCLs identified by the expression of TRAP (red) in the trabecular bone of the epiphysis. Upper left, higher magnification of the area indicated in the right panel showing the characteristic histology of the OCLs (arrow). Lower left, histologic image of a mouse tibia injected with MRTF-depleted D2.A1 cells showing a reduced number of OCLs in the epiphysis. Lower right, higher magnification of the area selected in the lower right image illustrating the characteristics of OCLs. Left panels: 40X, right panels: 200X, and insets: 1000X. **C)** Quantification of OCL numbers in mice injected with control vs MRTF-shRNA expressing D2.A1 (n=5 mice/group; each data point represents one mouse; *: p<0.05).

## DISCUSSION

In the present study, we report four major findings. First, while previous studies demonstrated MRTF-dependent changes in tumor-cell-intrinsic phenotypes (proliferation, migration and susceptibility to immune cell attack) (Medjkane *et al*., 2009; Tello-Lafoz *et al*., 2021; Gau *et al*., 2022), whether MRTF activity in tumor cells indirectly impacts other cells in the tumor microenvironment is not known. In this study, we uncover a novel tumor-cell-extrinsic function of MRTF in promoting osteoclastogenesis in a paracrine manner. Second, we establish CTGF as one of mediators of pro-osteoclastogenic paracrine action of MRTF in breast cancer cells. Although it is known that CTGF is one of the serum-induced early-responsive genes in cells (Herdegen and Leah, 1998), and its expression is induced in cultured cells when MRTF-A is overexpressed (Hinkel *et al*., 2014), CTGF is also regulated by other transcription factors (Cheng *et al*., 2018). Therefore, to what extent basal CTGF expression is dependent on MRTF’s action is not known. In this study, we show MRTF’s indispensable role in maintaining the basal CTGF expression in cultured cells, and a strong positive association between CTGF expression and MRTF-A gene signature in an actual human disease context. Third, we identify a host of additional cell-secreted immunoregulatory factors that are affected upon loss of MRTF expression in breast cancer cells providing additional mechanistic insights into how MRTF-deficiency in breast cancer cells could potentially interfere with osteoclastogenesis as depicted in **Fig 8**. Fourth, we demonstrate that MRTF-deficiency in breast cancer cells diminishes OCL abundance and bone colonization ability of tumor cells *in vivo*. Collectively, these findings suggest that MRTF inhibition could be an effective strategy to diminish OCL formation and skeletal involvement in breast cancer.

**Figure 8.**
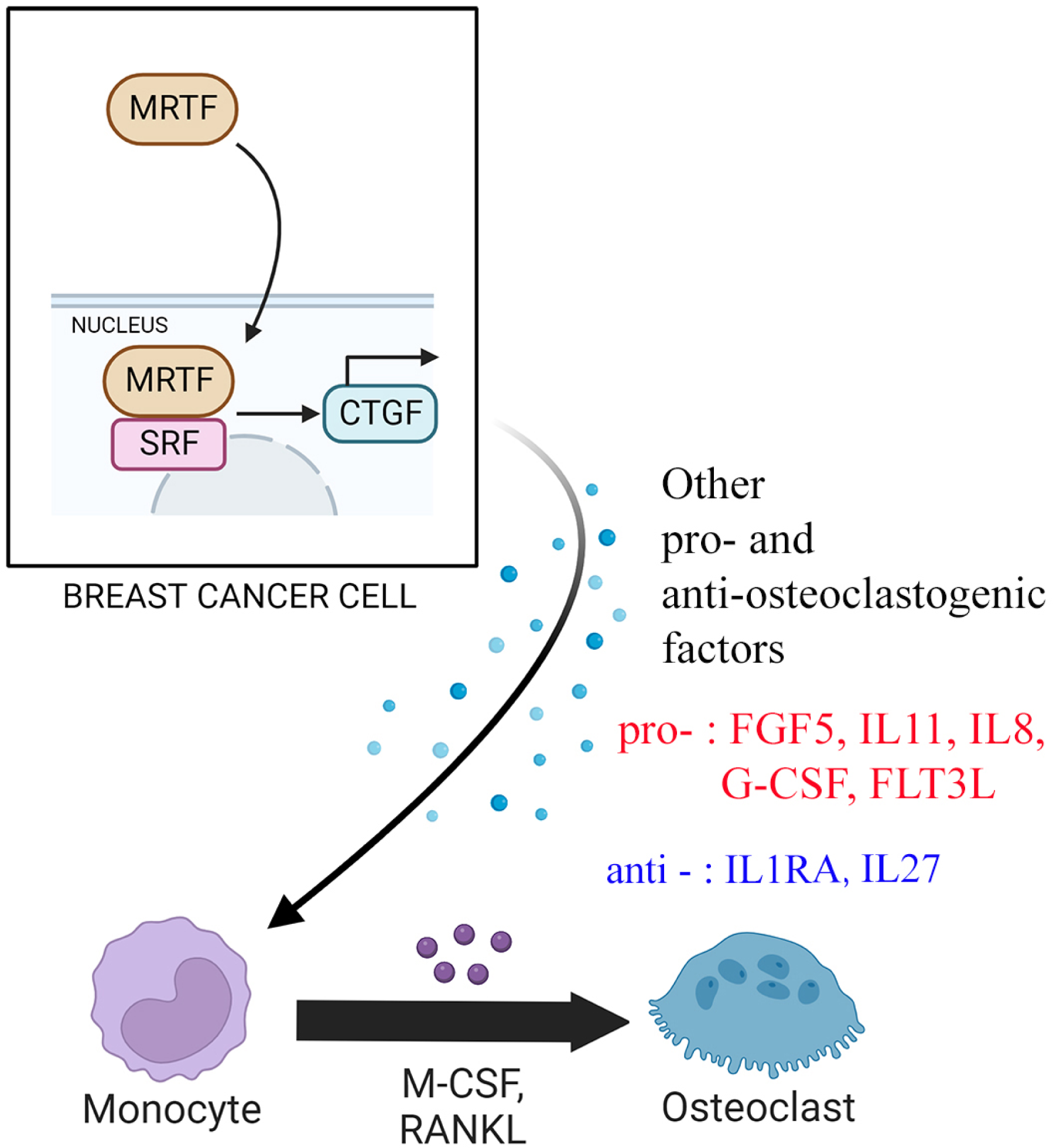
Proposed schematic model. MRTF activity in breast cancer cells promotes OCL differentiation through a paracrine action that involves the action of CTGF and other OCL-regulatory factors.

OCL differentiation and activation is a multistep process that is dependent on a multitude of growth factors liberated from the bone matrix and secreted by tumor cells in the metastatic niche. This involves proliferation and differentiation of hematopoietic stem cells to RANK-/TRAP-OCL progenitor cells, differentiation of OCL progenitor cells to RANK+/TRAP+ mononuclear OCL-precursor (a process that is dependent on the activities of NF-κb and NFATc transcription factors), and finally fusion of mononuclear OCLp cells to multi-nucleated and bone-resorption competent mature OCLs that are characterized by the expressions various markers including DC-Stamp, Atp6v0d2 (v-ATPase subunit d2), cathepsin K, and OSTM1 (osteoclastogenesis-associated transmembrane protein) (Amarasekara *et al*., 2018a). We revealed three key findings from our detailed analyses of TRAP-negative vs TRAP-positive (OCLp and OCL) cells in BMDM cultures. First, regardless of the MRTF expression status in breast cancer cells, exposure to tumor cell CM reduces the total cell count of BMDM cells (by ∼50%) but enhances their intrinsic OCL differentiation ability in an MRTF-dependent manner. We do not know which tumor cell secreted factor(s) is/are responsible for the anti-proliferative action on BMDMs. However, these results are also not totally surprising given that tumor cells secrete both pro-and anti-osteoclastogenic factors including those which suppress proliferation of myeloid-derived cells (IL27 as an example (Xu *et al*., 2024)) as also demonstrated by our Luminex data. Second, MRTF-proficient tumor cell-derived factors did not affect the total % of TRAP-positive cells (cumulative pool of OCLp cells and OCLs) in the BMDM culture but increased the OCL-to-OCLp ratio, a finding consistent with a scenario of accelerated transformation of OCLp to mature OCLs by tumor cell-secreted factors. Fusion of mono-nuclear OCLp is a critical process for generation of multinuclear OCLs. DC-Stamp, a transmembrane protein originally identified in dendritic cells, is essential for cell-cell fusion in OCLs (Yagi *et al*., 2005; Yagi *et al*., 2006). Our studies showed that RANKL-induced DC-Stamp expression in BMDMs is prominently enhanced (by ∼3 fold) by secreted factors of breast cancer cells in an MRTF-dependent manner. CTGF plays a key role in both induction of and interaction with DC- Stamp resulting in multi-nuclear OCL formation (Nishida *et al*., 2011). Since MRTF is critical for maintaining the basal expression of CTGF, MRTF’s requirement for increased OCL differentiation efficiency of BMDMs elicited by tumor cell-derived factors may be partly due to enhanced fusion of mono-nuclear cells by a CTGF/DC-Stamp signaling axis. Since cell motility plays a key role in bringing mononuclear OCLs into close proximity aiding their fusion process (Søe, 2020), another possibility is that BMDM motility is impacted by exposure to tumor cell secreted factors in an MRTF-dependent manner. Third, since the % of TRAP-positive BMDMs is drastically diminished in the presence of CM from MRTF-depleted tumor cells, we think that the early aspects of RANKL-induced differentiation process are somehow impaired by the paracrine effect of secreted factors of MRTF-depleted tumor cells. Future studies probing into downstream signaling readouts of RANKL such as activation and sustenance of NF-Kb signaling, as well as expression of early-responsive genes (e.g. NFATc1) will provide additional insights.

Our studies clearly indicate that MRTF-depletion in breast cancer cells has a suppressive effect on OCL differentiation process through a paracrine action that extends beyond the effect of CTGF. To this end, RNA-seq analysis identified MRTF-dependent upregulation of pro-osteoclastogenic factors IL11 and FGF5 (these factors were not represented in Luminex assay). Our Luminex studies additionally discovered that MRTF-depletion in breast cancer cells leads to reduced secreted levels of several pro-osteoclastogenic cytokines (IL8, G-CSF, IL25, FLT3L, IL1α) with concomitant upregulation of anti-osteoclastogenic cytokines (IL1RA, IL27, GM-CSF and RANTES/CCL5). Between these factors, G-CSF downregulation upon MRTF knockdown was reproducible in both human and mouse breast cancer cells. Among the pro-osteoclastogenic factors identified by Luminex analyses, IL8 stimulates OCL differentiation through inducing RANK-induced NFATc1 activation in an autocrine-signaling manner (Kopesky *et al*., 2014). G-CSF accelerates osteoblast turnover and suppresses the expression OPG (a negative regulator of RANK/RANKL signaling) (Christopher and Link, 2008) while eliciting increased OCL activity and bone resorption in both cancer (Hirbe *et al*., 2007) and periodontitis models (Yu *et al*., 2021). Although FL3T3L can substitute for M-CSF in supporting OCL induction (Lean, Fuller, and Chambers, 2001), since our studies were performed in cultures supplemented with M-CSF, we presume that reduced FL3TL secretion, if at all, has minimal role, in suppression of OCL differentiation in MRTF knockdown setting. Among the anti-osteoclastogenic factors, IL27 is known to inhibit RANKL-induced signaling pathway (Furukawa *et al*., 2009). IL1RA (IL1 receptor antagonist) inhibits the signaling elicited by IL1, an inducer of RANKL expression as well as an activator of various hallmark gene expression programs [NFκβ, p38, ERK] required for OCL differentiation (Kim *et al*., 2009)). Therefore, we will need to investigate potential contributions of these select candidate differentially abundant cytokines and changes in associated signaling pathways as additional future directions of mechanistic investigations underlying MRTF-regulated paracrine control of OCL differentiation. Although beyond the scope of the present study, given the concordance of downregulation of G-CSF and possibly IL1 signaling (marked by either downregulation of IL1α or upregulation of IL1RA) between human and mouse breast cancer cell lines, it will be important to design combinatorial rescue studies to determine whether exogenous supplementation of IL1 and/or G-CSF with CTGF fully recoups the OCL differentiation defect induced by MRTF-depletion in breast cancer cells. It is worth noting that many of the osteoclastogeneis-regulatory factors identified by our Luminex data were not found in the differentially expressed gene set in MDA-231 cells upon MRTF knockdown (Medjkane *et al*., 2009), possibly suggesting that MRTF depletion somehow affects secretion of these factors. Actin cytoskeleton plays an important role in exocytosis, and MRTF transcriptionally regulates components of actomyosin contractility (such as actin, MYL6, MYL9), mechanoresponsive proteins (integrins, YAP etc.) (Medjkane *et al*., 2009; Eder *et al*., 2024) and promotes cell contractility (Parreno *et al*., 2017) However, since our Luminex data showed evidence for both increase and decrease of secreted levels of OCL-regulatory factors, we speculate that it is unlikely that these changes are reflective of a global secretion defect of MRTF knockdown cells.

We previously showed that *in vivo* administration of CCG-1423, a small molecule that is widely used to inhibit MRTF’s function (Hayashi *et al*., 2014; Kobayashi *et al*., 2019; Song *et al*., 2022), dramatically reduces development of skeletal metastases following systemic inoculation of breast cancer cells (Gau *et al*., 2022). However, CCG-1423 or its derivative compounds have poly-pharmacology and are known to target other proteins (example: MICAL and pirin) (Lundquist *et al*., 2014; Lisabeth *et al*., 2019). In fact, CCG compounds bind to pirin at a much higher affinity than to MRTF. Importantly, a recent study shows evidence for global suppression of RNA synthesis by CCG compounds that extend far beyond MRTF’s action (Prajapati *et al*., 2024). Furthermore, in an *in vivo* administration setting, CCG compound’s action is targeted globally in a non-cell type-specific manner. Therefore, to what extent MRTF-activity in tumor cells contributes to bone colonization remained unclear in that study, which is now resolved by genetic proof-of-concept for MRTF’s role in bone colonization of breast cancer cells in this study. Since in this study, our experimental metastasis assessments were restricted to a relatively early time-point after tumor cell inoculation, reduced incidence of bone colonization of MRTF-knockdown cells is most likely caused by their defect in proliferation (as suggested by our proliferation assay data). However, our main finding that loss-of-function of MRTF in tumor cells negatively impacts OCL differentiation (this is further corroborated by our in vivo data) is still significant because of its implication in osteolysis-induced feed-forward loop to drive tumor cell growth in the bone microenvironment particularly in the advanced stage of metastatic growth of cancer cells.

Finally, we acknowledge that there are limitations of the present study. For example, this study models only the unidirectional signaling from cancer cells to OCLs and does not represent the complex multicellular crosstalk between osteoblasts, cancer cells and OCLs of the actual tumor microenvironment. Second, we did not investigate whether MRTF has any role in influencing the intrinsic OCL differentiation ability of BMDMs. Addressing these lines of questions using advanced model systems incorporating multiple cell types and further in vivo validation should be the focus for future investigations.

## EXPERIMENTAL PROCEDURES

### Animal Studies

All animal experiments were conducted in compliance with an approved IACUC protocol, according to University of Pittsburgh Division of Laboratory Animal Resources guidelines. Briefly, 10^5^ MDA-231 cells, suspended in 100 μL PBS, were injected into the left cardiac ventricle of fully anesthetized 5-week-old female NSG mice. For intratibial injections, 2.5x10^4^ D2.A1 cells, suspended in 20 μL PBS, were injected into the right tibia of fully anesthetized 5-week-old female immunocompetent Balb/c mice. Bioluminescence imaging (BLI) of anesthetized mice was performed with IVIS Spectrum 15 minutes after intraperitoneal injection of D-luciferin.

### Cell culture and OCL differentiation assay

Generation and culture of variants of MDA-231, stably co-expressing GFP and luciferase, and various MRTF-A constructs have been previously described (Gau *et al*., 2022). D2.A1 cells stably expressing luciferase (a gift from Dr. William Schiemann, Case Western Reserve University) were cultured in DMEM media (Lonza, catalog# BW12-604F) supplemented with 10% (v/v) FBS and 1% antibiotics (100 U/ml penicillin and 100 μg/mL streptomycin). For stable silencing of MRTF isoforms, MDA-231 and D2A1 cells were transduced with lentivirus encoding either MRTF-A/B shRNA (Vectorbuilder vector ID - VB200825-1246dwj) or non-targeting control shRNA (Vectorbuilder vector ID - VB151023-10034) and selected with puromycin. For transient transfection experiments, cells were transfected with smart-pool MRTF-A- and MRTF-B-specific siRNAs with smart-pool control siRNA (source catalog) serving as control as we previously described (Gau *et al*., 2022; Eder *et al*., 2024). Primary BMDMs were isolated from the long bones of female FVB mice and cultured in αMEM medium supplemented with 10% (v/v) fetal bovine serum (FBS; Corning, catalog# MT35011CV) and 1% antibiotics

(100 U/ml penicillin and 100 μg/ml streptomycin; Thermo Fisher, catalog# 15070063) as described previously (Allen-Gondringer *et al*., 2023). Primary BMDMs were seeded in the wells of a 96 wells plate in triplicate and expanded in αMEM supplemented with M-CSF (20 ng/mL) for 3 days to generate OCL progenitor cells. OCL differentiation was induced by additional 4 days of culture in the presence of RANKL (10 ng/mL) and M-CSF (10 ng/mL), with or without the addition of breast cancer cell CM at a 10% v/v ratio. To prepare CM, breast cancer cells were seeded in a 10 cm plate and cultured in serum containing media for 4 days prior to collection. For CTGF rescue studies, CTGF (PeproTech, 120-19) was added to the BMDM culture at either 50 ng/ml or 100 ng/ml concentration at the time of RANKL stimulation and continued until the end of experiments. To assess OCL differentiation, cells were fixed with 37% formaldehyde for 15 minutes, washed with PBS and stained using leukocyte acid phosphatase kit (Sigma-Aldrich, 387A-1KT) following the manufacturer’s instructions. OCLs (marked by TRAP+ multinucleated cells with three or more nuclei) were counted using the “OC_Finder” program which uses an automated cell segmentation approach before applying deep learning to classify the cells as either OCLs or non-OCLs (Wang *et al*., 2022). Total number of cells was counted using ImageJ.

### *In vivo* TRAP staining

The femur and tibia were dissected from mice intratibially injected with D2A1 cells. Samples were fixed in 4% paraformaldehyde for 24 hours post dissection and decalcified in 10% EDTA for 30 days. Slides were stained using leukocyte acid phosphatase kit (Sigma-Aldrich, 387A-1KT) following the manufacturer’s instructions. Image acquisition and quantification of OCLs were performed using an Axiostar plus microscope (Zeiss) and the AxioVision image analyzer software, respectively.

### 2D Proliferation Assay and protein quantification

Twenty-five thousand MDA-231 or D2A1 cells were seeded in the wells of a 6-well plate and cultured in Phenol free DMEM media (ThermoFisher, catalog# 31053028) supplemented with 10% (v/v) FBS and 1% antibiotics (100 U/ml penicillin and 100 μg/mL streptomycin). Cells were imaged and counted using Millicell DCI Digital Cell Imager four consecutive days starting from the day after cell seeding. Media samples were also taken daily and quantified for protein content using bicinchoninic acid assay.

### Calvaria preparation/tumor cell outgrowth assay

Female C57Bl/6 post-natal day 21 mice were injected intraperitoneally with 20 mg/kg Alizarin complex one dihydrate (CAS 455303-00-1, 2.0 mg/ml in bicarbonate buffer) and sacrificed 24 hours post injection. Calvaria were immediately excised and placed in phosphate buffered saline. The periosteum was removed by scraping with a scalpel. Calvaria were then digested in 0.5 mg/mL collagenase (Sigma Aldrich, catalog #C9697-50), 0.25% trypsin-EDTA for 10 minutes at a time at 37°C and stored in PBS at -20°C until used. Two thousand cells were seeded in duplicates in a 96-well plate on approximately 2 mm x 2 mm sized squares of calvaria. After 72 hours of culture, cells were imaged at 4x magnification and outgrowth was quantified by the total calvarial area covered by GFP+ cells using ImageJ.

### Quantitative RT-PCR

Total RNA was extracted from 2D cell cultures using RNeasy mini kit (Qiagen, catalog #74104) according to the manufacturer’s instructions. Complementary DNA (cDNA) was synthesized from 1 ug of RNA using the QuantiTect reverse transcription kit (Qiagen, catalog #205311) following the manufacturer’s instructions. Each PCR was prepared with 50 ng of cDNA, 12.5 μL of SYBR Select Master Mix (ThermoFisher, catalog #4472903), 1 μM forward and reverse primers, and water for a total volume of 25 μL. Thermal cycling and data analysis were performed using the StepOne Plus Real-Time PCR System and StepOne Software (Applied Biosystems) to detect quantitative mRNA expression of Cathepsin-K and DC-STAMP (primer sequences are listed in **supplementary Table S3**).

### RNA-sequencing and gene signature analyses

Differential gene expression data of MDA-231 with various MRTF perturbations are deposited in GEO (GSE253047). RNA-sequencing data processing/analyses and gene signature score analyses of MRTF-A by GSVA are detailed in our recently published study (Eder *et al*., 2024).

### Protein extraction and Western blotting

Cell lysates were prepared by a modified RIPA buffer (25 mM Tris–HCL: pH 7.5, 150 mM NaCl, 1% (v/v) NP-40, 0.3% SDS, 5% (v/v) glycerol), 1 mM EDTA, 50 mM NaF, 1 mM sodium pervanadate, and protease inhibitors (Sigma, catalog P8340) supplemented with 6× sample buffer diluted to 1× with the final SDS concentration in the lysis buffer equivalent to 2%. Conditions for the various antibodies were: polyclonal MRTF-A (Cell Signaling, catalog #14760S; 1:1000), MRTF-B (Cell Signaling, catalog #14613S; 1:1000), polyclonal GAPDH (Sigma, catalog #G9545, 1:2000), polyclonal CTGF (Abcam, catalog #ab6992), monoclonal Tubulin (Sigma, T9026, 1:1000). monoclonal anti-rabbit HRP (Jackson Immunoresearch, catalog #211–032-171, 1:1000), and polyclonal anti-mouse HRP (BD Biosciences, catalog #554002, 1:1000).

### Cytokine Analyses

CM harvested from 10 cm culture plates and concentrated to 100 µL was probed for cytokine/chemokine expression levels by HD48A (48-plex human discovery assay) and MD32 (32-plex mouse discovery assay) Luminex panels using a service provided by Eve Technologies (Calgary, AB, Canada). Values were normalized to the average value of the control group per analyte basis.

### Transwell migration assay

MDA-231 cells were serum-starved and seeded on the wells in triplicate in the wells of 24-well transwell chamber with 8 μm pores in serum-free media (#3422, Corning, USA) and allowed to migrate toward a chemotactic gradient of 10% serum in the culture media in the bottom chamber for either 6 hrs (for overexpression studies) or overnight (for knockdown studies). Non-migrated cells on the top of the well were removed using a cotton swab, and the bottom side of the membranes were fixed in 3.7% paraformaldehyde for 15 minutes and stained with DAPI to score the number of transmigrated cells using images taken with a 4X objective and analyzed by ImageJ software.

### Cell adhesion assay

The wells of a 96-well cell culture plate were coated with calcium phosphate (CaP) following a previously published protocol (Cen *et al*., 2022). CaP-coated wells were dried and pre-adsorbed with serum-containing media for 1 hour before seeding 10,000 MDA-231 cells in triplicates. After allowing cells to adhere for 4 hours at 37°C, non-adhered cells were removed by aspiration. Adherent cells were then fixed with 3.7% formaldehyde for 15 minutes at room temperature and stained with DAPI to visualize nuclei. Images were acquired using Lionheart automated imaging system at 4X magnification, and cell quantification was performed using ImageJ software.

### F-actin staining

Cells were stained with rhodamine-conjugated phalloidin following previously described protocol (Gau *et al*., 2022; Eder *et al*., 2024). The average rhodamine fluorescence in GFP+ cancer cells (acquired with 20X objective) was quantified for comparison between the control and the experimental groups.

### Statistics

All statistical tests were performed using GraphPad Prism 9 software. ANOVA (for more than 2 groups) and Student’s t-test (for 2 groups) were performed for comparing the means between various experimental groups. For ANOVA, we performed Tukey’s post hoc test for individual group comparisons. A p-value less than 0.05 was considered statistically significant.

## DATA AVAILABILITY

RNA-sequencing datasets are available in GEO (GSE253047).

## SUPPORTING INFORMATION

The article contains supporting information.

## ACKNOWLDEGMENTS

DG was supported by a National Cancer Center fellowship, NCI K99-CA267180 and T32-HL129964. IE was supported by NIBIB T32-EB001026 and NCATS TL1-TR001858. VY was supported by a summer undergraduate research fellowship from the School of Engineering, University of Pittsburgh. Fangyuan Chen was a former visiting research scholar at the University of Pittsburgh School of Medicine supported by funds from The China Scholarship Council and Tsinghua University. Research in the Roy lab is supported by grants from the NIH (R01CA248873, R01CA271095, and R21EY032632) and the Department of Defense (HT9425-24-1-0556).

## AUTHOR CONTRIBUTION

PC, DG, IS and NW performed experiments, analyzed data, and wrote manuscript; VY performed experiments and analyzed data, DB, NW, IE, and FC performed data analyses, JT contributed to generation of experimental tools; AVL oversaw bioinformatics data analyses, AL assisted with osteoclast detection and quantification in bone histology slides, DLG supervised some aspects of the project, PR conceived the overall study design, supervised the project, oversaw experiments, edited/wrote manuscript and acquired funding.

## List of abbreviations (in alphabetical order)

BLI: Bioluminescence imaging
BMDM: Bone marrow derived macrophages
CM: Conditioned media
CTGF: Connective tissue growth factor
DC-Stamp: Dendritic cell-specific transmembrane protein
Dox: Doxycycline
ECM: Extracellular matrix
ER: Estrogen receptor
G-CSF: Granulocyte colony stimulating factor
GSS: Gene signature score
GSVA: Gene signature variance analyses
HER: Human epidermal growth factor receptor
IL: Interleukin
OB: Osteoblast
OCL: Osteoclast
OPG: Osteoprotegerin
MCSF: Macrophage colony-stimulating factor
PTHrP: Parathyroid hormone-related protein
MKL: Megakaryoblastic leukemia
MRTF: Myocardin-related transcription factor
NFκB: Nuclear factor kappa B
OX: Overexpression
PR: Progesterone receptor
SAP: SAF-A/B, acinus, PIAS
SRF: Serum-response factor
RANKL: Receptor activator for nuclear factor κB
TCGA: The Cancer Genome Atlas
TNBC: Triple-negative breast cancer
TNF: Tumor necrosis factor
TRAP: Tartrate resistant alkaline phosphatase
WT: Wild-type

