## Supplementary Info for "Breast cancer cells promote osteoclast differentiation in an MRTF-dependent paracrine manner"

(Chawla et al.)

#### Chawla et al. Figure S1

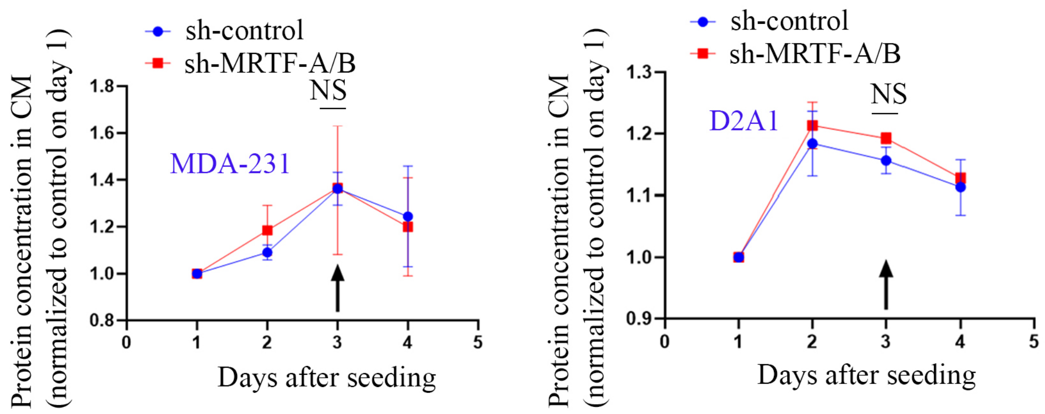


**Figure S1: MRTF knockdown does not affect the protein concentration in the CM of breast cancer cells.** Protein concentration measured in the CM of control vs stable MRTF-A/B knockdown MDA-231 and D2A1 cells in 2D tissue culture on different days after seeding (data normalized to the value measured for control groups of cells on day 1); arrow indicates the day CM was harvested for OCL differentiation assays).

#### Chawla et al. Figure S2

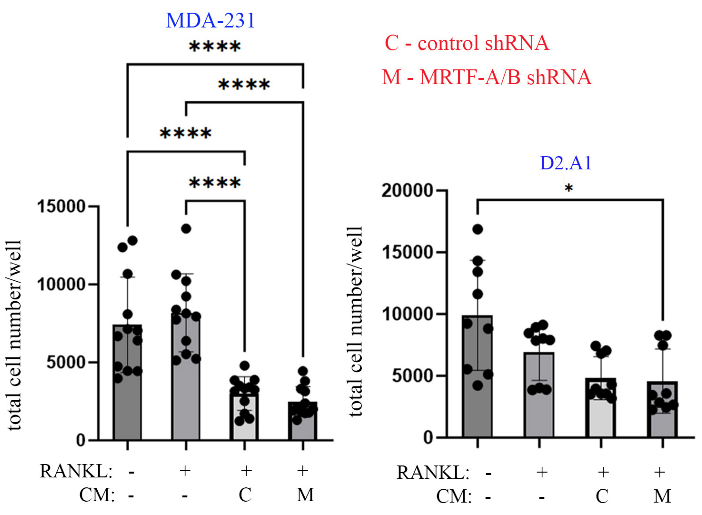


**Figure S2: Addition of tumor cell-derived factors reduce proliferation of BMDMs.** Quantifications of total cell count of BMDM cultures for the indicated treatment groups (data summarized from 4 and 3 independent experiments for CM treatment from MDA-231 and D2.A1, cells respectively (* p<0.05, **: p<0.01, and **** p<0.0001).

**Chawla et al. Figure S3**

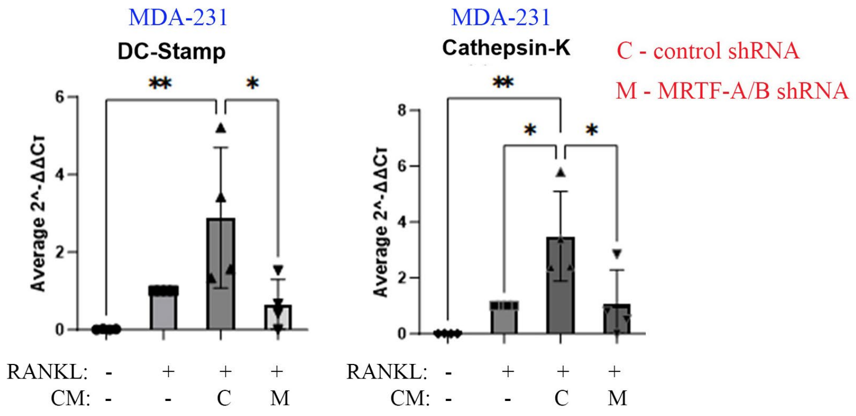


**Figure S3: Effect of tumor cell-derived factors on OCL-related gene expression in BMDMs.** Quantitative RT-qPCR analyses show the fold-changes in the mRNA expression levels of DC-Stamp and Cathepsin-K in BMDM cultures subjected to the indicated treatments (fold changes normalized to the expression values for RANKL-alone treatment group; data summarized from 3 independent experiments; * $p<0.05$, and **: $p<0.01$).

#### Chawla et al. Figure S4

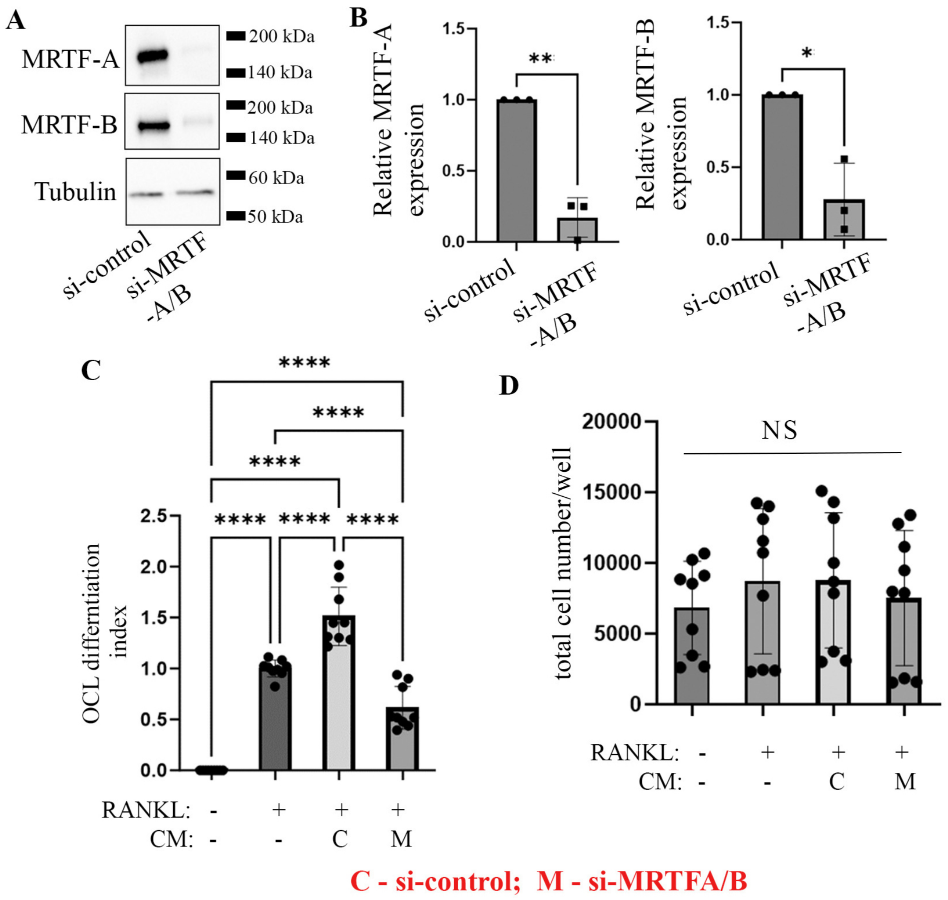


**Figure S4: Validation of the effect of MRTF depletion in breast cancer cells on paracrine modulation of OCL differentiation in a transient knockdown setting. A-B)** Representative immunoblots (*panel A*) and quantification of immunoblots (*panel B;* data summarized from 3 experiments) of MDA-231 lysates showing siRNA-mediated transient knockdown of MRTF isoforms by siRNAs (isoform-specific MRTF siRNAs were pooled for transfection). **C-D)** Summary of OCL differentiation index (*panel C*) and total cell count (*panel D*) of BMDM cultures subjected to the indicated treatments**;** data summarized from 3 independent experiments (*: $p<0.05$; **: $p<0.01$; ****: $p<0.0001$; NS – not significant).

**Chawla et al. Figure S5**

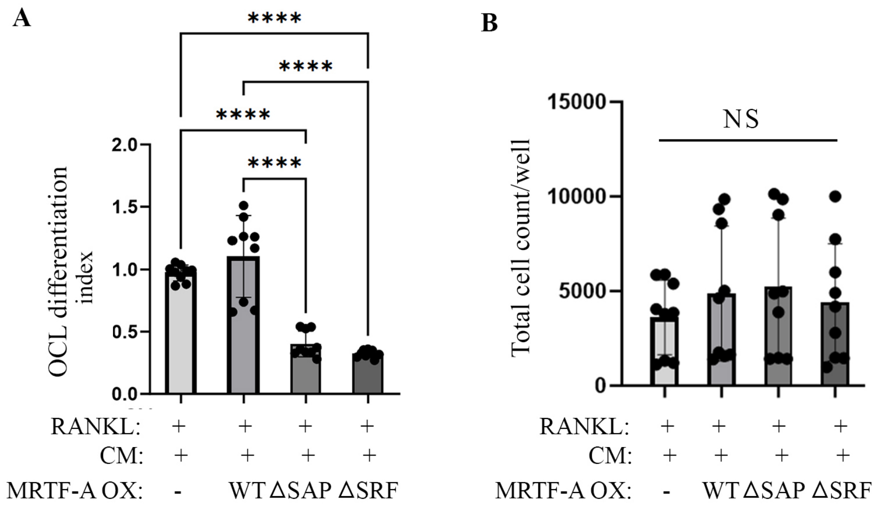


**Figure S5**: **Both SAP-domain and SRF-related functions of MRTF are important for tumor-cell-directed paracrine modulation of OCL differentiation.** Quantifications of OCL differentiation indices (*panel A*) and total cell count (*panel B*) of RANKL-stimulated BMDMs without or with supplementation of CM derived from the 2D cultures of control vs various stable MRTF-A (either WT or specific functional mutants) overexpressing sublines of MDA-231 cells (data summarized from 3 independent experiments, **** $p<0.0001$; N.S – not significant).

#### Chawla et al. Figure S6

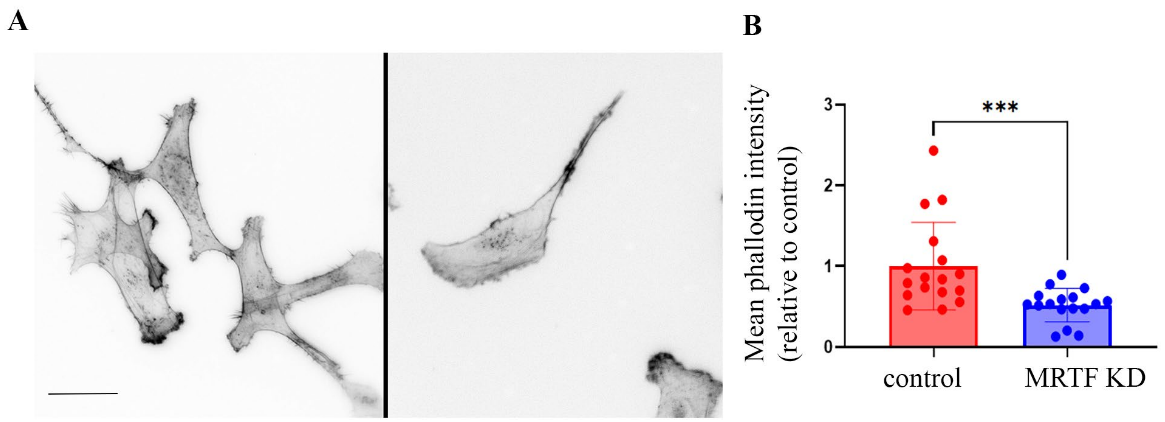


**Figure S6**: **MRTF depletion reduces overall F-actin content in MDA-231 cells.** Representative images (*panel A;* 40X micrographs) and quantification of average fluorescence intensity (*panel B*) of rhodamine-phalloidin stained control vs MRTF-A/B knockdown (MRTF KD) MDA-231 cells (data are summarized from 3 independent experiments with at least 45 cells analyzed per experiment per group based on images acquired with a 20X objective; ***: $p<0.001$; scale bar – 20 μm).

### Chawla et al. Figure S7

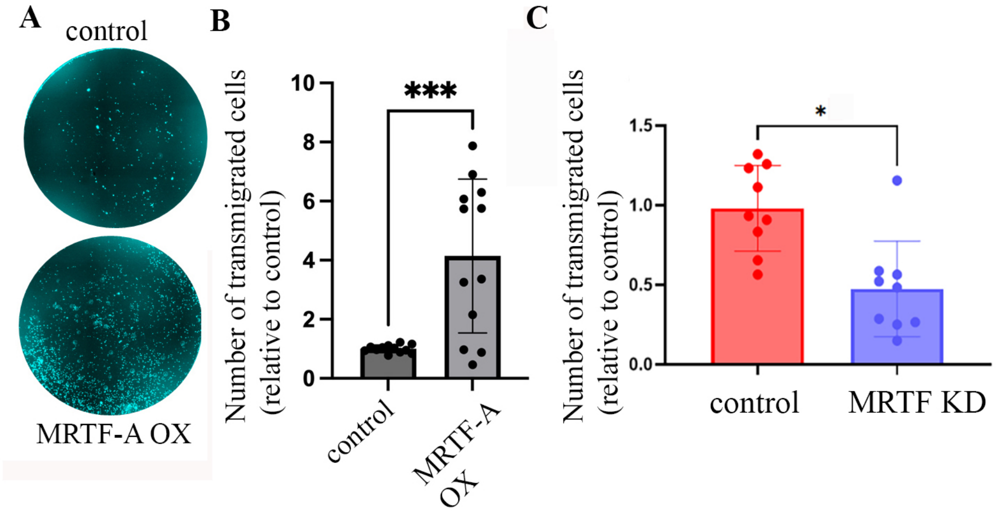


**Figure S7**: **MRTF promotes chemotactic migration of MDA-231 cells. (A-B)** Representative images of DAPI-stained transmigrated cells (*panel A*) and quantification (*panel B*) of relative migration of control vs MRTF-A overexpressing (MRTF-A OX) MDA-231 cells under a chemotactic gradient of 10% serum. **C)** Quantification of relative trans-migration of control vs MRTF-A/B knockdown (MRTF KD) MDA-231 cells. Data are summarized from 5 and 3 independent experiments in overexpression and knockdown settings, respectively; p<0.05; ***: p<0.001).

#### Chawla et al. Figure S8

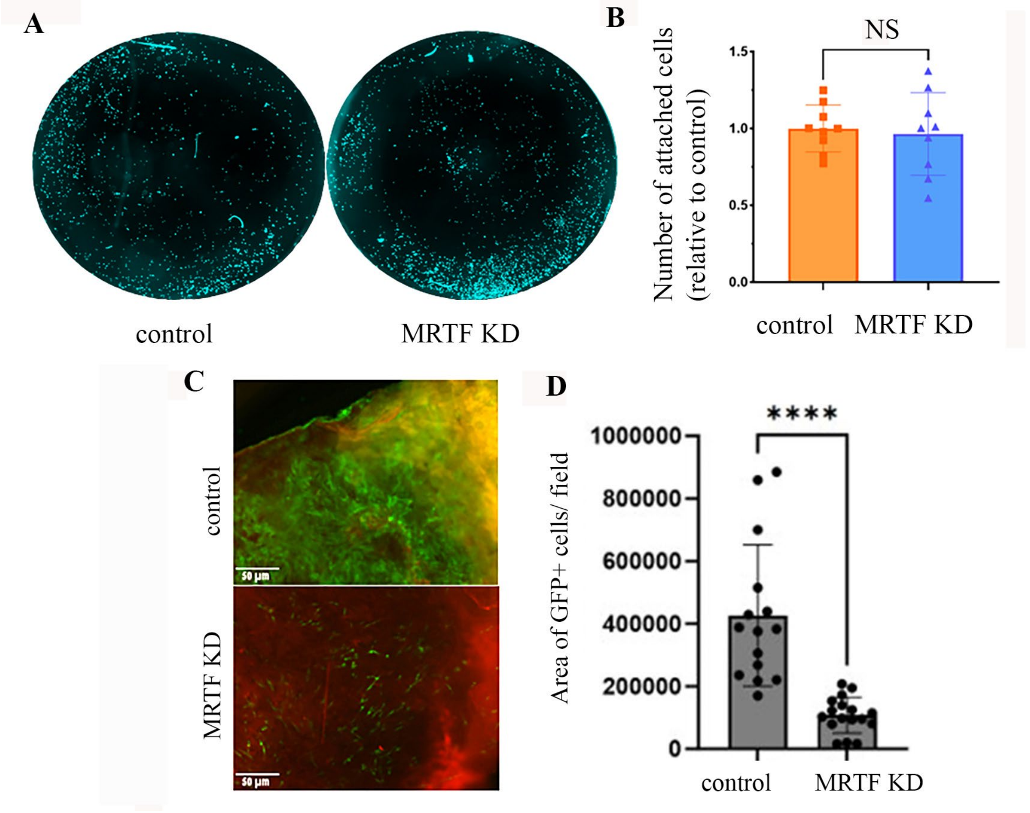


**Figure S8**: **Effect of MRTF knockdown on adhesion and outgrowth of MDA-231 cells. (A-B)** Representative images of DAPI-stained adherent cells (*panel A*) and quantification (*panel B*) of relative adherent control vs MRTF-A/B knockdown (MRTF KD) MDA-231 cells 4 hours after seeding on calcium-phosphate coated tissue-culture substrate (data are summarized from 3 independent experiments; NS – not significant). **C)** Representative fluorescence images of GFP+ control and MRTF KD MDA-231 cells 72 hrs after seeding on alizarin red+ mouse calvaria bone punches. **D)** Quantification of area of bone covered with GFP+ cells 72 hours post-seeding (data summarized from 3 independent experiments, **** $p<0.0001$; scale bar – 50 µm). Non-flat nature and autofluorescence of calvarial bone punches account for the sub-optimal quality of fluorescence images.

**Supplementary Table S1:** Breast cancer bone metastasis gene signature

| Direction of change | Genes |
|---|---|
| Upregulated | *CXCR4, MMP1, ADAMTS1, FGF5, CTGF, IL11, FST, NFAT1, Ph4, and BBS1* |
| Downregulated | *APOBEc3B, NUP155, MFAP3L, NIP7, C6orf17, KCNS1, STEAP3, C16orf61, PALM2, ATL2, and SFT2D2* |

** compiled from Kang, et al. 2003; Savci-Heijink, et al. 2016

**Supplementary Table S2**: List of cytokines/chemokines analyzed by Luminex assays

| Luminex Panel | Cytokines/Chemokines probed |
|---|---|
| *HD-48plex* (human) | sCD40L, EGF, Eotaxin, FGF-2, Flt-3 ligand, Fractalkine, G-CSF, GM-CSF, GROα, IFNα2, IFNγ, IL-1α, IL-1β, IL-1ra, IL-2, IL-3, IL-4, IL-5, IL-6, IL-7, IL-8, IL-9, IL-10, IL-12p40, IL-12p70, IL-13, IL-15, IL-17A, IL-17E/IL-25, IL-17F, IL-18, IL-22, IL-27, IP-10, MCP-1, MCP-3, M-CSF, MDC (CCL22), MIG, MIP-1α, MIP-1β, PDGF-AA, PDGF-AB/BB, RANTES, TGFα, TNFα, TNFβ, VEGF-A |
| *MD-32plex* (mouse) | Eotaxin, G-CSF, GM-CSF, IFNγ, IL-1α, IL-1β, IL-2, IL-3, IL-4, IL-5, IL-6, IL-7, IL-9, IL-10, IL-12p40, IL-12p70, IL-13, IL-15, IL-17A, IP-10, KC, LIF, LIX, MCP-1, M-CSF, MIG, MIP-1α, MIP-1β, MIP-2, RANTES, TNFα, VEGF-A |

**Supplementary Table S3**: RT-PCR Primer details

| Gene name | Primer sequence |
|---|---|
| Cathepsin-K | 5’-GAAGAAGACTCACCAGAAGCAG-3’ (sense)<br>5’-TCCAGGTTATGGGCAGAGATT-3’ (antisense). |
| DC-Stamp | 5'-TTGCCGCTGTGGACTATCTG-3' (sense)<br>5'-GAATGCAGCTCGGTTCAAAC-3' (antisense) |
